# Epicardial contributions to fibro-inflammatory signaling in a Pkp2-deficient arrhythmogenic cardiomyopathy model

**DOI:** 10.1101/2025.10.08.680826

**Authors:** Daniel D. Han, Alan C. Brooks, Cameron D. Baker, Ronald A. Dirkx, Deanne M. Mickelsen, Benjamin Fisler, Kavya Phadke, John M. Ashton, Mario Delmar, Eric M. Small

## Abstract

**Background:** Arrhythmogenic Cardiomyopathy (ACM) is an inherited disease that is characterized by lethal ventricular arrhythmias stemming from myocyte dysfunction. ACM is associated with considerable subepicardial fibrosis and inflammation with right ventricle predominance. Most cases of gene positive ACM are caused by a desmosome protein mutation, with plakophilin-2 (*Pkp2*) mutations being most common. We hypothesized *Pkp2*-deficiency in epicardium-derived cells (EPDCs) contributes to fibro-inflammatory signaling and ACM pathogenesis.

**Methods:** We developed transgenic mice that lack *Pkp2* in cardiomyocytes (Pkp2-cKO) or in both cardiomyocyte and EPDC (Pkp2-ceKO) via the tissue-specific expression of tamoxifen-inducible Cre recombinase. Non-myocyte populations were isolated 21 days post-tamoxifen injection for single cell RNA-sequencing (scRNA-seq). Immunohistochemistry, flow cytometry, qRT-PCR, and echocardiography were used to interrogate cardiac physiology and cellular composition.

**Results:** We identified a population of epicardium-derived fibroblasts characterized by the expression of *Ccl2*, *Ccl7*, *Thbs1*, and *Ptx3* that accumulated upon *Pkp2* deletion. While pro-inflammatory EPDCs are found in Pkp2-cKO mice, they become significantly enriched in Pkp2-ceKO mice. Pro-inflammatory fibroblasts acquired the senescence-associated secretory phenotype (SASP), correlating with elevated Senescence Associated (SA)-βgal staining in the right ventricle. Gene expression, flow cytometry, and histological data also revealed an exaggerated inflammatory response in Pkp2-ceKO mice, that progresses from right to bi-ventricular predominance. Importantly, macrophages and B cells accumulate in both Pkp2-cKO and Pkp2-ceKO mice compared to controls. Antibody-mediated B cell depletion delays the early inflammatory and fibrosis response but did not significantly alter end-stage cardiac physiology.

**Conclusion:** *Pkp2* deletion in EPDC facilitates the emergence of a fibro-inflammatory phenotype that may contribute to ACM pathogenesis.

## Introduction

Arrhythmogenic Cardiomyopathy (ACM), also known as the Arrhythmogenic Right Ventricular Cardiomyopathy/Dysplasia (ARVC/D), is an inherited cardiac disorder that primarily affects young adults.^1^ Most cases of gene positive ACM are caused by mutations in genes that encode desmosome proteins, structural components of the cardiomyocyte intercalated disk. Plakophilin-2 (*Pkp2*) mutations account for around 40% of the observed desmosome protein mutations in ACM patients. Mutations in other desmosome proteins including desmoglein-2 (*Dsg2*), desmocollin-2 (*Dsc2*), desmoplakin (*Dsp*), and plakoglobin (*Jup*) occur less frequently.^2,3^ The primary clinical manifestation of ACM is ventricular arrhythmia and ultimately sudden cardiac death (SCD) if untreated. ACM is the most common cause of SCD after hypertrophic cardiomyopathy among patients who are <35 years old.^4–5^ As many patients with ACM are asymptomatic, misdiagnosis of ACM is common and SCD often occurs before diagnosis.^6^

The most notable cardiac phenotype of ACM is the progressive buildup of fibrofatty deposits.^7^ Fibrofatty deposits initially form in the free wall of the right ventricle and progress into the left ventricle to cause biventricular fibrosis.^7^ Furthermore, clinical studies have revealed that fibrofatty deposits are initially observed in the sub-epicardium, expanding later into the myocardium towards the endocardium.^8,9^ The epicardium is a source of progenitor cells that differentiate into various non-myocyte cardiac lineages during embryonic development, including cardiac fibroblasts, smooth muscle cells, and pericytes.^10,11^ The epicardium also becomes re-activated after myocardial insult, and expresses pro-inflammatory paracrine signals, suggesting the epicardium and its descendants may play a role in cardiac inflammation and remodeling after injury.^12,13^

*Pkp2* deficiency alters the epicardial phenotype in vitro by reducing cell adhesions and stimulating epicardial to mesenchymal transition.^14,15^ However, evidence that *Pkp2*-deficient epicardium-derived cells (EPDCs) contribute to ACM pathogenesis has not been demonstrated in vivo. Therefore, we generated mice lacking *Pkp2* in cardiomyocytes (Pkp2-cKO) or in both cardiomyocytes and EPDCs (Pkp2-ceKO) to investigate potential contributions of *Pkp2*-deficient EPDCs in ACM pathogenesis. Using single-cell RNA sequencing (scRNA-seq) we defined the emergence of a pro-inflammatory fibroblast population, particularly in Pkp2-ceKO mice. The fibro-inflammatory program originating from EPDCs correlates with cellular senescence and the induction of the senescence associated secretory phenotype (SASP). We further found Pkp2-ceKO mice exhibit an overabundance of macrophages, B cells, and T cells in the heart, progressing from RV to LV predominance. While ablating B cells in mice using a blocking antibody significantly delayed the development of fibrosis, this approach did not alter end stage cardiac physiology. Taken together, our results establish the role of EPDCs in the emergence of a senescent and pro-inflammatory phenotype that may contribute to ACM disease pathogenesis.

## Methods

All methods and key materials can be found in Supplemental Methods and the Major Resources Table in the Supplemental Material. scRNA-sequencing data has been deposited in the Gene Expression Omnibus (GEO) database under accession code GSE300141.

## Results

### Epicardium-endocardium gradient of fibro-inflammatory deposition following *Pkp2* deletion

Using a previously developed mouse line that exhibits many of the clinical characteristics of ACM following cardiomyocyte (CM)-specific *Pkp2* deletion (called Pkp2-cKO) ^16^, we observed fibrotic foci originating at the epicardial surface (Supplemental Figure 1A). To investigate the impact of *Pkp2* deletion in EPDCs on fibrosis, we generated mouse lines harboring tamoxifen-inducible Cre recombinase under the control of various combinations of *Wt1* and *Myh6* responsive transgenes (Wt1-CreERT2^17^ and Myh6-MerCreMer^18^), to cause recombination of floxed *Pkp2* alleles in the epicardium and cardiomyocytes, respectively (Figure 1A, B). These mice also harbor a *Rosa^mTmG^* allele to label the recombined cells with green fluorescence protein (GFP). Picro-Sirius red stained heart sections showed comparable biventricular fibrosis in Pkp2-ceKO mice and Pkp2-cKO at 21 days post tamoxifen injection (dpi) (Figure 1C), which was significantly elevated compared to controls and Pkp2-eKO (epicardium deletion alone) mice (Figure 1C). Serial echocardiography also revealed comparable decline in cardiac physiology in Pkp2-ceKO and Pkp2-cKO mice, compared to controls at 21 and 42 dpi, which progresses from RV to bi-ventricular predominance (Supplemental Figure 1B). These findings reveal that both Pkp2-cKO and Pkp2-ceKO mice capture the fibrotic ACM pathophysiology.

**Figure 1.**
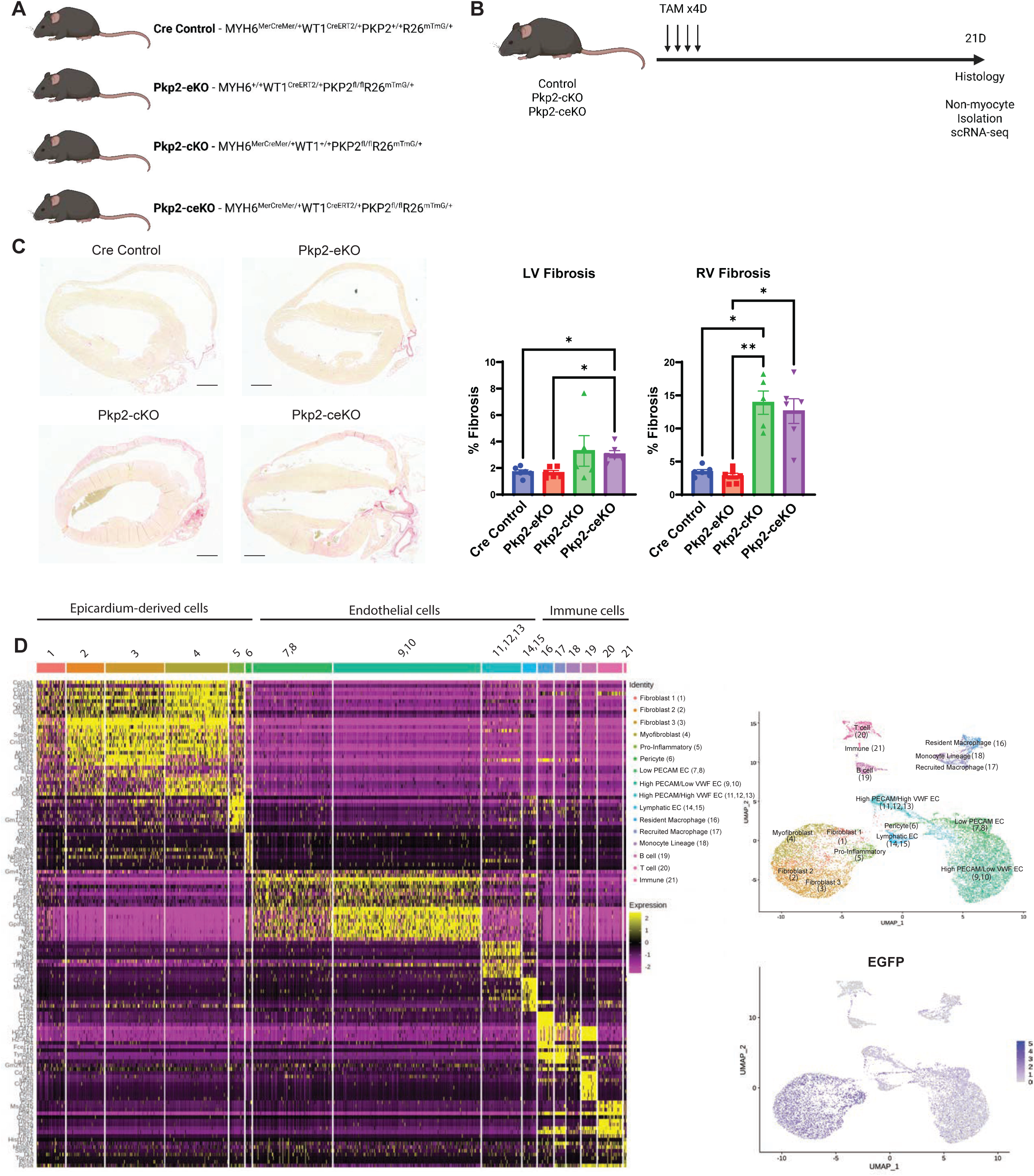
Defining the single cell transcriptome of non-myocytes in a Pkp2 deficient mouse model of ACM. **A)** Schematic describing Pkp2 gene deletion strategy in cardiomyocytes, epicardial cells, or both cell-types, compared to control. **B)** Schematic describing experimental timeline for tamoxifen injections, histology and scRNA-seq. **C)** Picrosirius Red stained heart sections (left) and quantification (right) reveals fibrosis in the left ventricle (LV) and right ventricle (RV) in the various genotypes. Quantification of LV and RV fibrosis represented by mean +/- SD. Scale bar=1mm. **D)** Hierarchical clustering (left) and uniform manifold approximation and projection (UMAP, right top) defines non-myocyte identities from Cre Control, Pkp2-cKO and Pkp2-ceKO mice. Data is obtained from cells pooled from n=2 hearts per genotype. Epicardium-derived cells were identified as GFP expressing cells (right bottom).

### Single cell transcriptomics identifies pro-inflammatory fibroblasts in Pkp2-ceKO mice

Since EPDCs give rise to fibroblasts and *Pkp2* is expressed in the epicardium^14^, we probed whether *Pkp2* deletion in EPDCs alters their transcriptional phenotype. Hearts from mice of genotypes described above were digested via Langendorff Perfusion at 21 dpi and the non-myocyte fraction was subjected to single cell RNA sequencing (sc-RNAseq) using the 10X Genomics platform. We observed comparable RNA features, counts, and percent mitochondrial between the various genotypes (Supplementary Figure 2). Hierarchical clustering and uniform manifold approximation and projection (UMAP) captured the transcriptional signature of fibroblasts, endothelial cells, mural cells, and immune cells (Figure 1D). EPDCs were identified based on expression of *eGFP,* revealing a majority of EPDCs are mesenchymal with minimal contributions to endothelial cell populations (Figure 1D, Supplemental Figure 3A, B).

UMAP revealed the expression of *Tcf21* and *Pdgfrα* across all cell populations to define epicardium-derived fibroblasts (Figure 2A). We defined 3 pro-fibrotic clusters (4, 5, and 6) based on high expression of *Postn* and various collagens and a reduction in *Tcf21* expression (Figure 2A, B). Cluster 4 contains activated fibroblasts based on co-expression of *Meox1*^19^ (Figure 2A), while cluster 6 is composed of activated pericytes based on the lack of *Meox1* and the robust expression of *Pdgfrb* and *Cspg4*^20^ (Figure 2B). Notably, cluster 5 exhibits a distinct pro-inflammatory phenotype based on the expression of pro-inflammatory cytokines including *Ptx3*, *Ccl2*, *Ccl7*, *Thbs1*, and *Il6* (Figure 2B). When pseudobulk analysis of sc-RNAseq was performed across all the genotypes, we observed an inverse relationship between pro-inflammatory and fibrosis marker genes at 21dpi, with Pkp2-ceKO mice exhibiting higher levels of pro-inflammatory cytokines and lower levels of fibrosis markers, compared to Pkp2-cKO mice (Figure 2C, D). Indeed, Pathway Enrichment analysis revealed an over-representation of genes regulated by transcription factors related to immune pathways in Pkp2-ceKO such as *Rela, Relb, Stat3, Brd4, Runx1,* and *Runx2* in Pkp2-ceKO non-myocytes (Supplemental Figure 3C). Importantly, overlaying genotype onto UMAP space in conjunction with proportionality analyses correlated the emergence of a robust pro-inflammatory fibroblast phenotype in Pkp2-ceKO mice with a significant immune cell burden (Figure 2E). Indeed, raw counts reveal a near complete absence of pro-inflammatory fibroblasts in control mice, which expand to become a major component of the non-myocyte fraction in Pkp2-ceKO mice.

**Figure 2.**
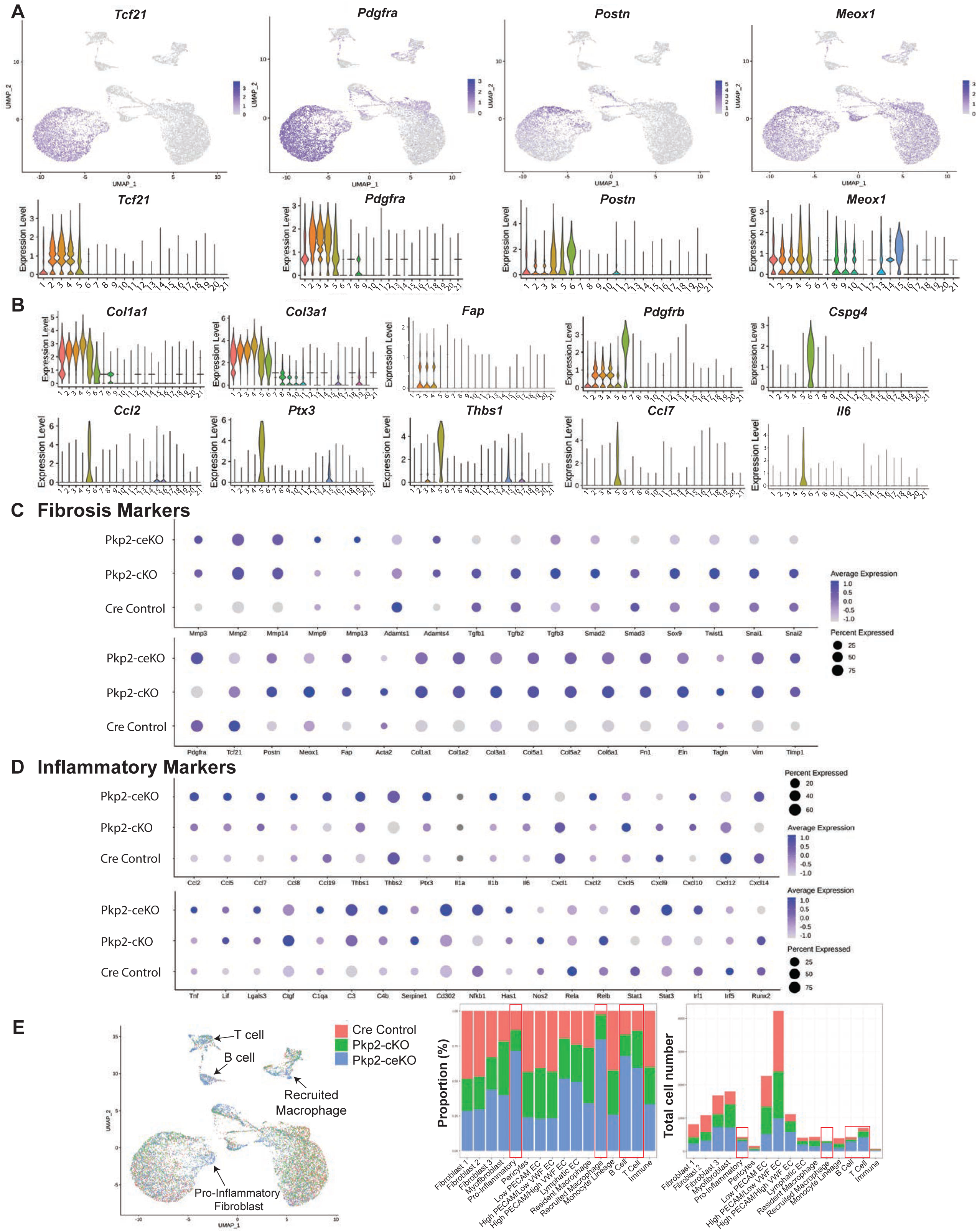
scRNA-seq reveals excessive fibro-inflammatory signaling in Pkp2-ceKO mice. **A)** UMAP (top) and violin plots (bottom) characterize the expression of fibroblast markers (*Tcf21, Postn, Meox1, Pdgfra*) B) Violin plots reveal activated fibroblasts (*Col1a1, Col3a1, Fap*), pericytes (*Pdgfrb, Cspg4*) and pro-inflammatory fibroblasts (*Ccl2, Ptx3, Thbs1, Ccl7, Il6*). **C, D)** Pseudobulk analysis reveals the relative expression of various markers of fibrosis (C) and inflammation (D) in the various genotypes. E) UMAP (left) and proportionality histograms (right) reveal the distribution of cell types among different genotypes.

Sub-analysis of EPDCs revealed 9 populations with unique transcriptional signatures (Figure 3A, B). *Tcf21* and *Pdgfrα* defined all fibroblasts, and *Postn, Meox1, Fap,* and *Acta2* defined activated fibroblasts (Fig. 3C, D). This analysis also better resolved the pro-inflammatory fibroblast population (Cluster 7), which is significantly enriched in Pkp2-ceKO mice, apparent in Pkp2-cKO, and nearly non-existent in controls (Figure 3B). Pro-inflammatory fibroblasts uniquely express *Ccl2, Ptx3, Ccl7, Thbs1*, and *Il6* (Figure 3E). Pseudotime trajectory analysis revealed the most naïve (quiescent) fibroblast population as cluster 3, which expresses the most *Tcf21*. Lineage trajectory 2 terminates with cluster 7 pro-inflammatory fibroblasts, defined by gene ontology terms related to inflammatory signaling and apoptosis (Figure 3F). Additional pseudotime trajectories yielded quiescent fibroblasts (Cluster 0, Lineage 1), activated fibroblasts (Cluster 5, Lineage 3), and pericytes (Cluster 8, Lineage 4) (Supplemental Figure 4). Consistent with the emergence of a pro-inflammatory fibroblast phenotype, cytokine array revealed an elevation in IFN-γ, Serpine1, Ptx3, M-CSF, and Postn in peripheral blood of Pkp2-cKO mice at 21 dpi compared to controls (Figure 3G). Since most of the cytokine signals are likely to be locally produced in the damaged myocardium, our data suggests a systemic pro-inflammatory signal may originate from epicardium-derived fibroblasts.

**Figure 3.**
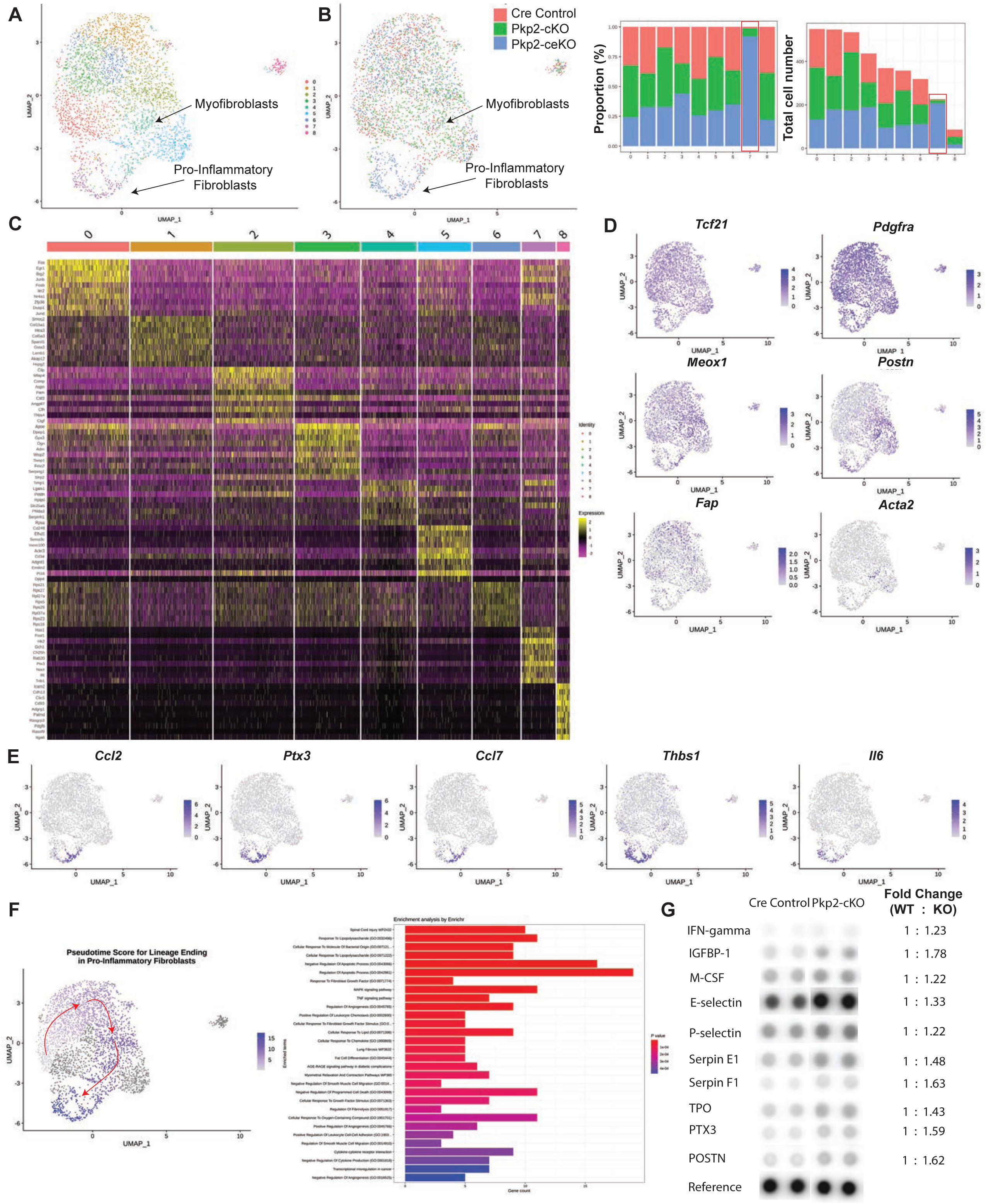
Sub-analysis of epicardium-derived cells reveals emergence of a pro-inflammatory fibroblast upon Pkp2 deletion. **A)** UMAP defines the identity of epicardium-derived non-myocyte lineages. **B)** UMAP (left) and proportionality histograms (right) define the relative distribution of cell phenotypes among different genotypes. Cluster 7 is significantly enriched in Pkp2-ceKO mice. **C)** Heirarchical clustering based on the most differentially expressed genes defines nine epicardium-derived mesenchymal sub-types. **D)** UMAP defines the distribution of all fibroblasts (*Pdgfra Tcf21*) and activated fibroblasts within cluster 4 (*Meox1, Postn, Fap, Acta2*). **E)** UMAP establishes localization of pro-inflammatory markers (*Ccl2, Ptx3, Ccl7, Thbs1, Il6*) in cluster 7, which is enriched in Pkp2-ceKO mice. **F)** Pseudotime trajectory (left) and Enrichr analysis (right) reveals a distinct lineage terminating in fibroblast cluster 7 that is characterized by gene ontology terms related to inflammation. **G)** Serum was obtained from control and Pkp2-cKO mice at 21 dpi. Cytokine Array reveals elevated levels of pro-inflammatory cytokines in Pkp2-cKO. Data represents serum pooled from n=3 (controls) and n=4 (Pkp2-cKO).

### Pkp2 deletion induces senescence and the Senescence-Associated Secretory Phenotype (SASP) in non-myocytes

The senescence-associated secretory program (SASP) refers to secreted factors expressed by senescent cells that contribute to negative paracrine and systemic effects^21^. Senescence associated (SA)-βgal staining revealed an accumulation of senescent cells in the RV (but not LV) of Pkp2-cKO mice at 21 dpi; no positive stain was observed in controls (Figure 4A). Given *Ccl2, Thbs1, Ptx3, Ccl7, Il6* are highly expressed in Pkp2-ce and -cKO mice, we utilized a published panel of SASP factors to identify whether the pro-inflammatory fibroblast population may express markers of cellular senescence^21^. The expression of *Apoe* and *Tmem176b* is detected in many non-myocytes, along with *Cdkn1a, Mt1*, and *Mt2* (Figure 4B). Among EPDCs, the most robust SASP signature is in pro-inflammatory fibroblasts (Figure 4C and Supplemental Figure 5). Pseudobulk analysis of EPDCs from all genotypes revealed the enrichment of SASP and p53-dependent cellular senescence in Pkp2-ceKO mice (Figure 4D). It is important to note that Pkp2-ceKO mice exhibit higher expression of DNA damage markers (*Atm, Atr, Hmga2*), suggesting DNA damage may initiate p53-mediated senescence (Figure 4C).

**Figure 4.**
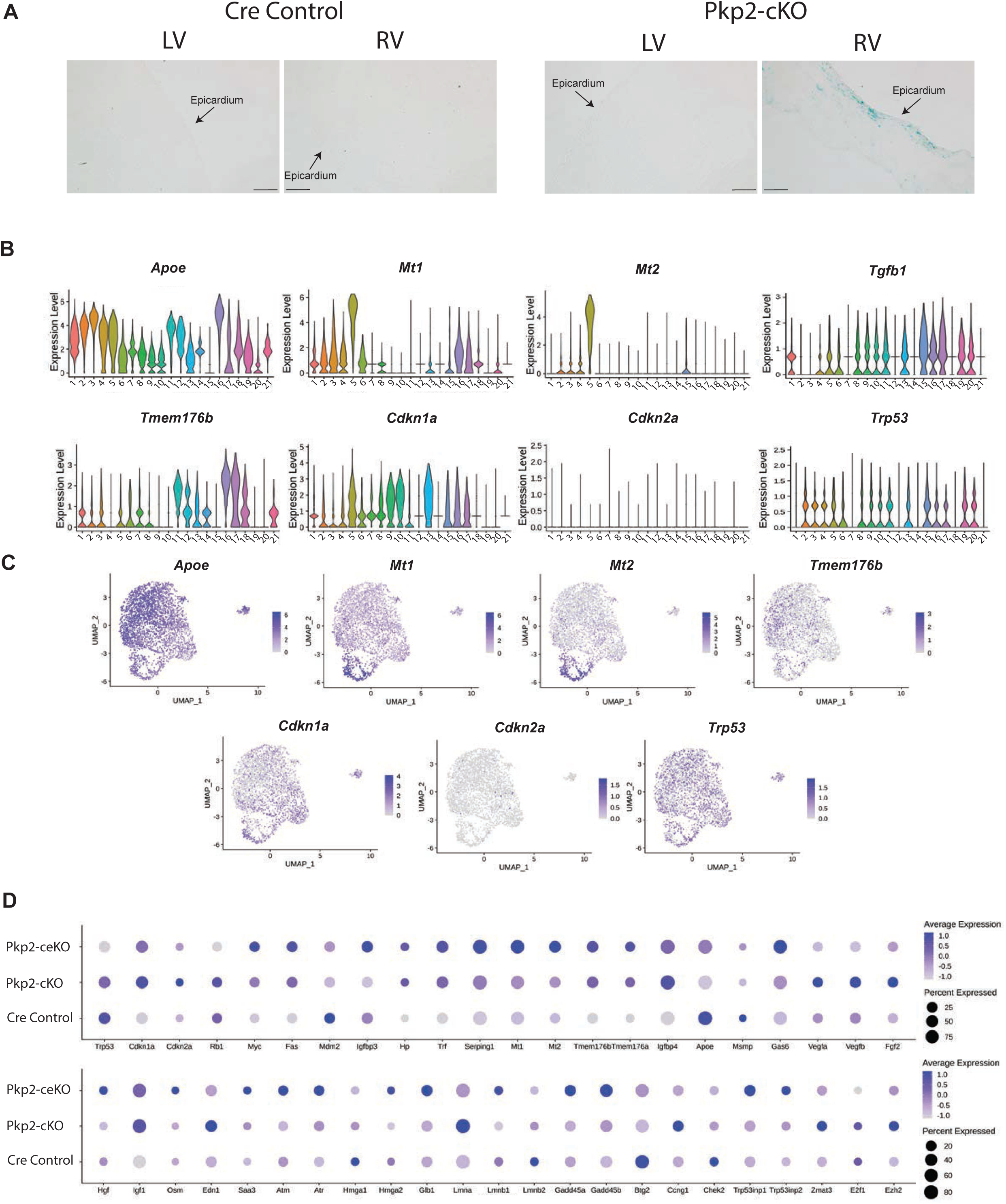
Pro-inflammatory fibroblasts arising in Pkp2-cKO mice exhibit signs of cellular senescence. **A)** Heart sections obtained from control and Pkp2-cKO hearts at 21 dpi were stained with SA -βgal as a marker of cellular senescence. Scale bar=100mm. **B, C)** Violin plots (B) and UMAP (C) of non-myocyte scRNA-seq data reveal cellular distribution of senescence and Senescence-Associated Secretory Phenotype (SASP) markers. **D)** Pseudobulk analysis reveals relative expression of senescence and SASP markers between genotypes.

### Increased immune cell burden in the Pkp2-ceKO heart

Induction of the pro-inflammatory SASP in Pkp2-ceKO mice correlates with increased immune cell burden (See Figure 2E). Here, we used *Ptprc (Cd45)* expression in Violin plots and UMAP to confirm the identity of all macrophages, B cells, and T cells (Figure 5A). High expression of *Adgre1 (F4/80)* and *Cd68* define cluster 16, 17, and 18 as macrophages and monocytes (Figure 5B). Cluster 16, which is identified in equivalent numbers in Pkp2 mutant and control genotypes, contains resident macrophages based on high *Cx3cr1* and low *Ccr2* expression^22,23^ (Figure 5B). Additional resident macrophage markers, *Mrc1, Cd163, Ccl24,* and *C1qA* were also enriched in cluster 16^24^ (Figure 5C and Supplemental Figure 6A). In contrast, recruited macrophages that highly express *Ccr2* (Cluster 17) are more abundant in Pkp2-ceKO hearts (Figure 5C and see Figure. 2E). Recruited macrophages in cluster 17 also express *Plac8, Ly6c2, Illb,* (Figure 5C and Supplementary Figure 5). The pseudobulk analysis of macrophage clusters between all genotypes confirmed macrophage burden in Pkp2-ceKO mice with higher gene expression of *Cd68, Ly6c2, Ccr2, Spp1, Tgfb1, Il1b, Il6, Lgals3, Plac8*^22^ (Figure 5D). Finally, we identified the Pkp2-ceKO enriched cluster 19 as B cells based on expression of *Cd19*, *Cd79a*, *Cd79b,* and other B cell markers (Figure 5D, E and Supplementary Figure 6B); while Cluster 20 contains T cells based on high expression of *Cd3e*, *Cd4*, *Cd8a*, *Ifng, Ccr7,* and other T cell markers (Figure 5D, E and Supplementary Figure 6C).

**Figure 5.**
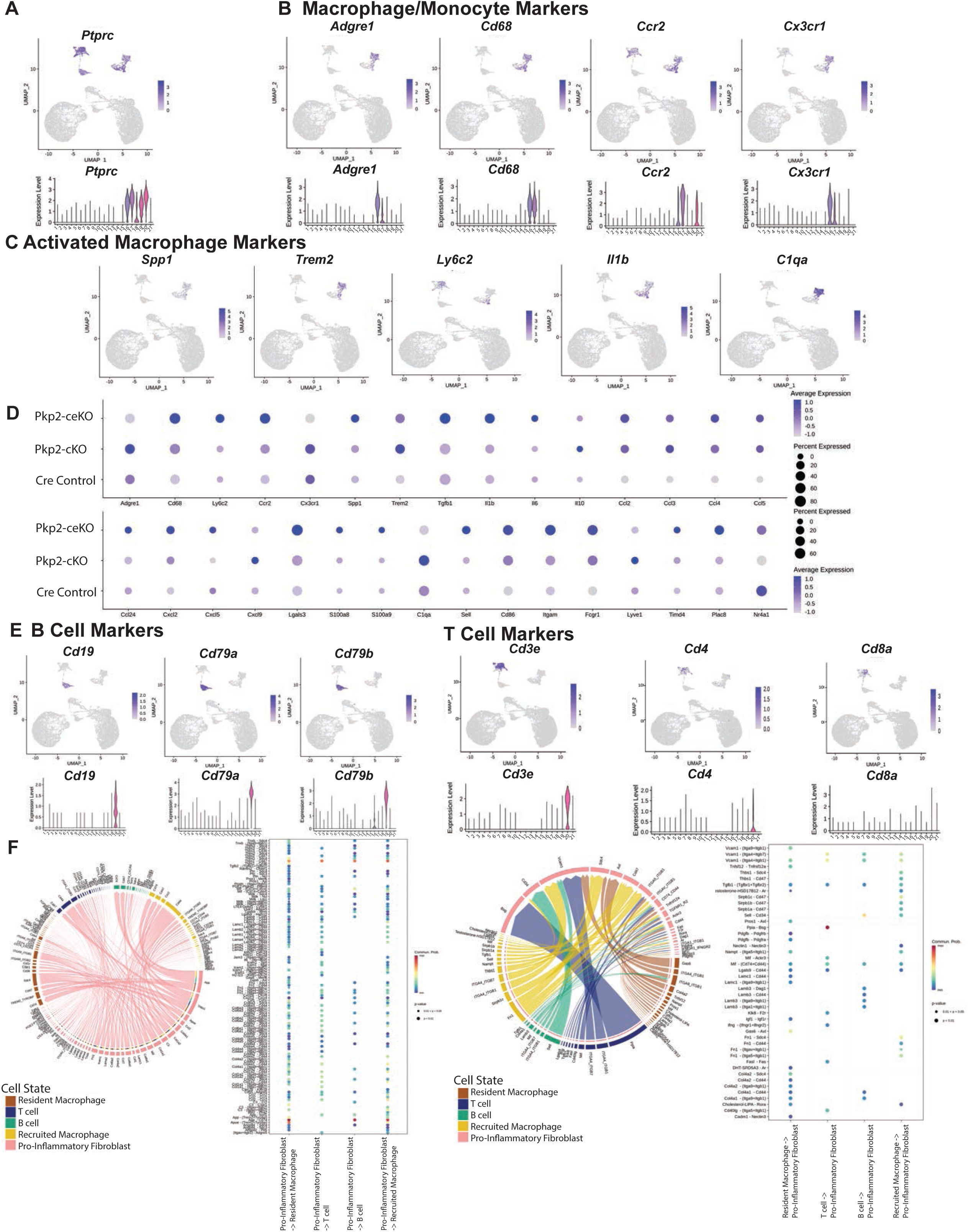
Immune cell recruitment to the Pkp2 deficient heart. **A)** Localization of *Ptprc* in UMAP (top) and violin plots (bottom) from non-myocyte scRNA-seq reveals identity of hematopoietic cells. **B)** Cell clusters defined by general macrophage/monocyte markers in UMAP (top) and violin plots (bottom). **C)** Cell clusters defined by activated macrophage markers in UMAP plots. D) Pseudobulk analysis of non-myocyte scRNA-seq data reveals relative expression of immune cell markers between all genotypes. E) Cell clusters defined by B cell (left) and T cell (right) markers in UMAP (top) and violin plots (bottom). F) CellChat defines intercellular communication from pro-inflammatory fibroblasts to immune cells (left) and from immune cells to pro-inflammatory fibroblasts (right).

Given the abundance of both pro-inflammatory cytokines and immune cells in Pkp2-ceKO heart, we used CellChat to define interactions between all cell populations. This analysis identified immune cell recruitment/retention signals arising from pro-inflammatory EPDC cluster 11 (Figure 5F). The key pathways include Thsb1-CD47, which is present in all immune cells, in addition to Ccl2-Ccr2 and Ccl7-Ccr2 signaling, which are present in the resident and recruited macrophage and T cells. An additional unique feature is the C3-Cr2 pathway, which is biased towards B cells and is often involved in the autoimmunity. Indeed, a prior report described features of autoimmunity and autoantibody formation in ACM.^25^

To identify triggers of the pro-inflammatory phenotype, we performed CellChat between the pro-inflammatory fibroblasts and whole ventricle transcriptomes obtained via bulk RNA-seq comparing Pkp2-cKO hearts to control hearts^16^. However, this did not yield significant interactions other than collagen-related pathways (Supplemental Figure 7). Therefore, fibroblast activation may not be driven by factors displaying differential expression in cardiomyocytes at 21dpi, or may be driven by stress pathway that do not depend on alterations in gene expression such as damage associated molecular patterns (DAMPs).

### B cell depletion reduces early stage ventricular remodeling in Pkp2-ceKO mice

Based on flow cytometry, Pkp2-ceKO hearts displayed an increase in the CD45+ immune cell burden between 10 and 21 dpi (Figure 6A, B). Moreover, there was a trend towards increased B cell burden in Pkp2-ceKO mice at 21 dpi, although this did not reach statistical significance. (Figure 6B). The role of B cells in cardiac pathologies has gained considerable attention; however, conflicting reports suggest beneficial or detrimental effects of B cell in heart^26,27^. Therefore, we tested whether B cell depletion with anti-CD20 antibody infusion alters ACM pathogenesis in Pkp2-ceKO mice (Figure 6C). Flow cytometry confirmed B cell depletion following anti-CD20 infusion compared to IgG isotype control at 7 and 21 dpi (Figure 6D). Although the RV diameter trended smaller in the Pkp2-ceKO infused with anti-CD20, we did not observe a significant difference in any functional parameter based on echocardiography at 21 dpi (Figure 6E). In contrast, B-cell depletion elicited a significant decrease in the amount of LV fibrosis at 21 dpi (Figure 6F).

**Figure 6.**
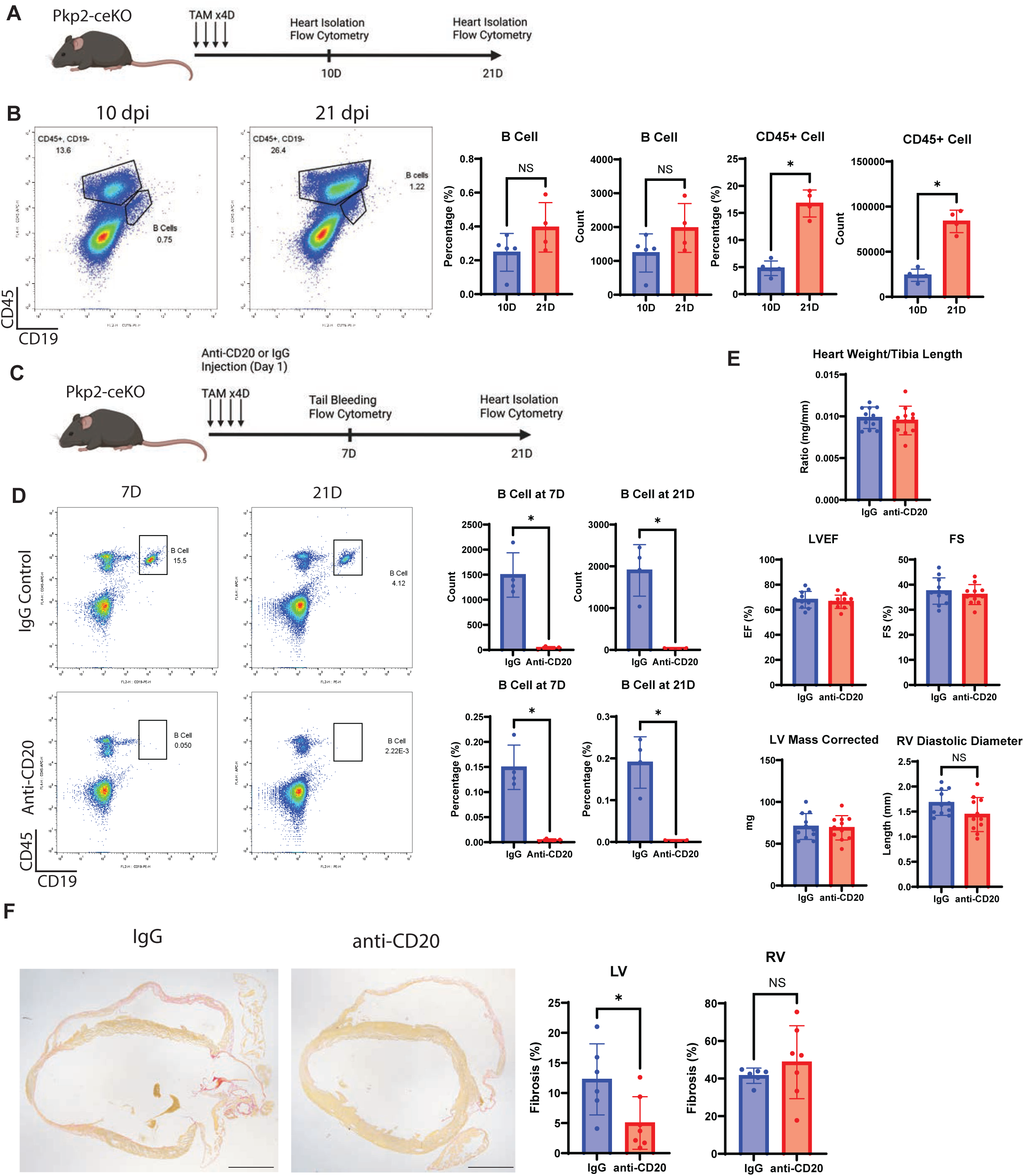
B cell depletion delays early inflammatory and fibrotic responses without improving end stage ACM pathophysiology. **A)** Schematic describing experimental timeline of cardiac immune cell quantification. **B)** Heart cells obtained from Pkp2-ceKO hearts at 10 dpi and 21 dpi were subjected to flow cytometry (left) to define all hematopoietic cells (CD45+) and B cells (CD19+). Quantification of flow cytometry data represented by average +/- SD (right). Data analyzed by Students t-test. \**P*<0.05. **C)** Schematic describing experimental timeline for B Cell Depletion and analysis. **D)** Flow cytometry performed on peripheral blood (7 dpi) and cardiac non-myocytes (21 dpi) to define all hematopoietic cells (CD45+) and B cells (CD19+). Quantification of flow cytometry data represented by average +/- SD (right). Data analyzed by Students t-test. *P<0.05. **E)** Cardiac physiology measurements obtained at 21dpi. Left ventricle ejection fraction (LVEF), Fractional Shortening (FS), LV Mass and RV diastolic diameter obtained by echocardiography. Data represented by mean +/- SD (right). Data analyzed by Students t-test. NS, not significant. F) Heart sections obtained at 21dpi from Pkp2-ceKO mice infused with isotype IgG control or anti-CD20 antibodies were stained with Picrosirius red to visualize fibrosis (left). Quantification of LV and RV fibrosis represented by mean +/- SD (right). Data analyzed by Students t-test. \**P*<0.05. Scale bar = 1mm.

We confirmed efficient Cre-dependent deletion of Pkp2 in both Pkp2-cKO and Pkp2-ceKO hearts by qRT-PCR (Figure 7A). The expression of the B cell marker *Cd19* is significantly elevated in the LV of Pkp2-ceKO mice treated with IgG control, compared to Pkp2-cKO mice, while anti-CD20 treatment decreased *Cd19* expression in the LV (Figure 7B). Fibrosis markers (*Postn*, *Col1a1*, *Col1a2*, and *Col3a1*) and pro-inflammatory markers (*Ccl2*, *Thbs1*, *Ptx3*, and *Ccl7*) are elevated in both the LV and RV of Pkp2-ceKO mice infused with IgG control; in contrast, Pkp2-cKO mice display an increase in these inflammatory and fibrosis markers primarily in the RV. B cell depletion led to a significant reduction in the expression of several fibrosis and inflammation markers in the LV of Pkp2-ceKO mice but did not reduce the fibrotic and inflammatory program in the RV (Figure 7C and 7D). Interestingly, senescence markers (*Cdkn1a*, *Apoe*, and *Tmem176b*) were also enriched in hearts obtained from Pkp2-cKO and Pkp2-ceKO mice infused with IgG control. *Cdkn1a* elevation in the LV was suppressed by anti-CD20 treatment, while *Apoe* and *Tmem176b* were only elevated in the RV and not impacted by B cell depletion (Figure 7E).

**Figure 7.**
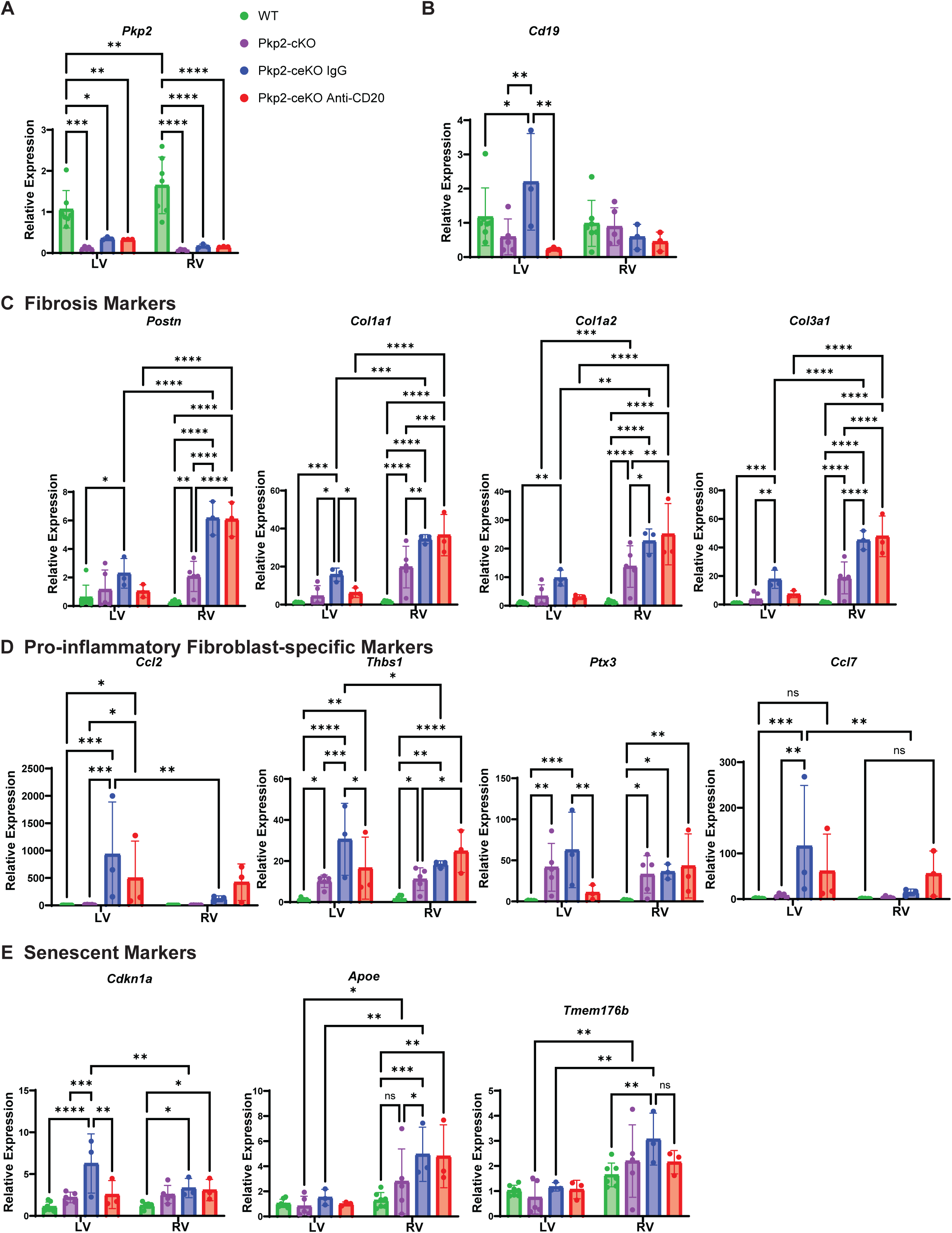
Cardiac chamber specific changes in fibro-inflammatory program upon Pkp2 deletion and B-cell depletion. The entire LV and RV freewall was obtained from various treatment groups at 21dpi for qRT-PCR to define gene expression changes. **A)** *Pkp2* expression confirms tamoxifen mediated deletion. B) *Cd19* expression represents B cell abundance. C) Expression of activated fibroblast markers and various collagens to define fibrosis phenotype. D) Expression of pro-inflammatory genes representing cluster 11 fibroblasts. D) Expression of markers of cellular senescence. Data represented by average +/- SD. Data analyzed by Two-way ANOVA with Tukey post-hoc test. \**P*<0.05 \*\**P*<0.01, \*\*\**P*<0.001, \*\*\*\**P*<0.0001. ns, not significant.

At 21 dpi in our Pkp2 deletion model, the disease has progressed to the point of significant RV deterioration, while the LV exhibits early signs of cardiac pathology. qRT-PCR highlights the abundant expression of recruited immune cell markers such as *Adgre1, Ccr2, Cx3cr1, Spp1,* and *Trem2* in the RV compared to LV (Figure 8A, B and see Figure 7D). In contrast, genes encoding secreted pro-inflammatory factors such as *Ccl2*, *Ccl7, Ptx3,* and *Thbs1* are enriched in the LV compared to the RV (Figure 8B and see Fig. 7D). This data reinforces the concept that the pro-inflammatory phenotype is a transient and early response that precedes the recruitment of inflammatory cells, demonstrated here as a wave from the RV to the LV. The prevention of LV fibrosis and inflammation with anti-CD20 infusion suggests B cells may regulate early inflammatory and fibrotic responses captured by LV dysfunction at 21 dpi; however, its effects may eventually be masked by macrophage and T cell burden during the later stages of ACM pathogenesis, which is captured here by RV remodeling .^28^

**Figure 8.**
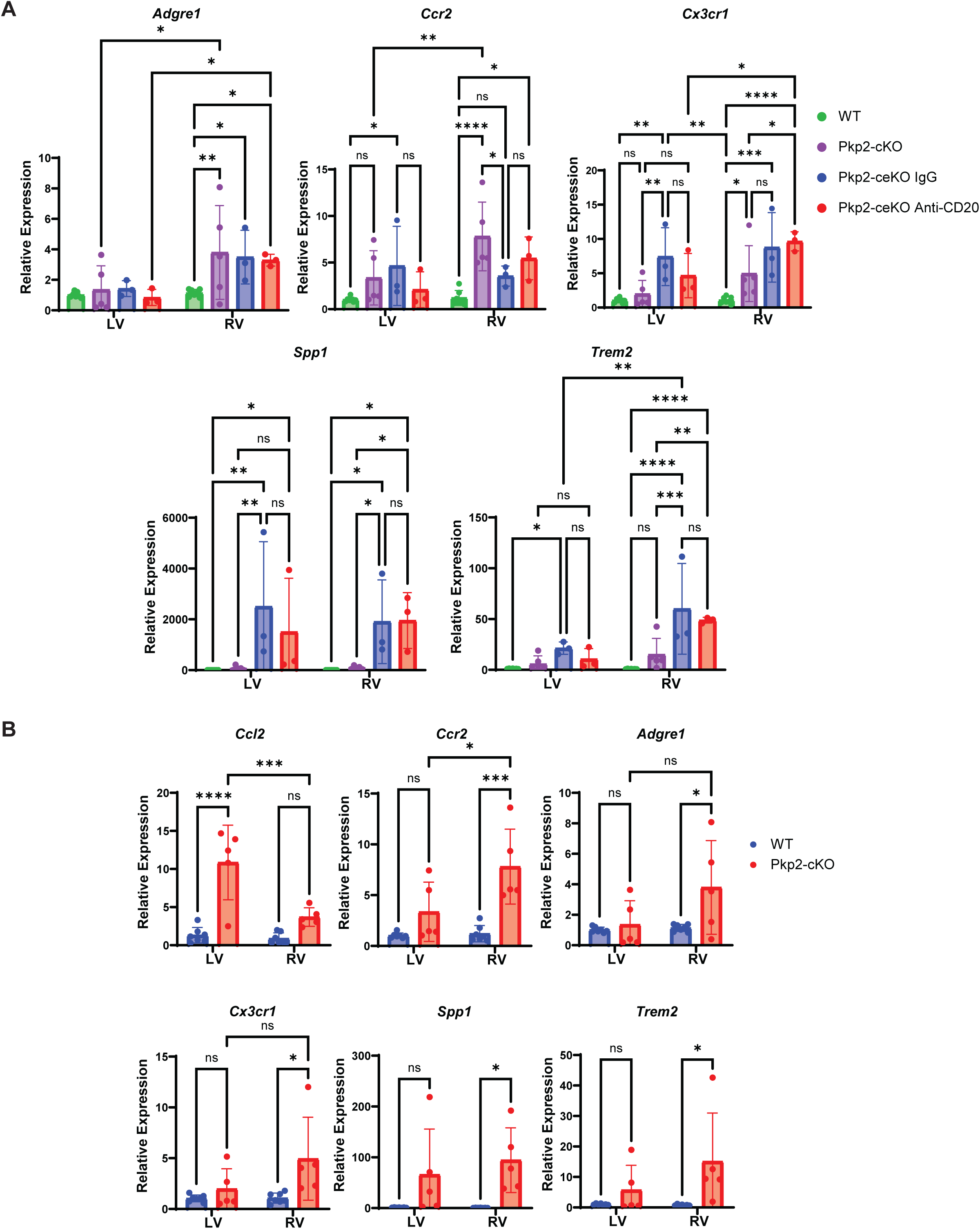
Temporal progression of inflammatory response from RV to LV revealed by qRT-PCR. The entire LV and RV freewall was obtained from various treatment groups at 21dpi for qRT-PCR to define gene expression changes. A) Macrophage marker gene expression. B) Macrophage and pro-inflammatory cytokine genes exhibit inverse expression in LV compared to RV. Data represented by average+/- SD. Data analyzed by Two-way ANOVA with Tukey post hoc test. \**P*<0.05 \*\**P*<0.01, \*\*\**P*<0.001, \*\*\*\**P*<0.0001. ns, not significant.

## Discussion

In this study, we identified a population of senescent and pro-inflammatory fibroblasts that emerge following deletion of *Pkp2* in cardiomyocytes and epicardial cells. In addition, we observed an accumulation of macrophages, T cells, and B Cells in Pkp2-ceKO mice. While B cell depletion did not improve end stage cardiac physiology or RV fibrosis following Pkp2 deletion, it did ameliorate LV fibrosis, suggesting a potential early therapeutic window to rescue LV function in ACM. Taken together, this study defines a novel role for non-myocytes in amplifying inflammatory and senescent signaling and immune cell recruitment in Pkp2-deficient ACM, which regulates fibrotic burden.

While ACM disease pathology initiates in the sub-epicardium of the RV^7–9^, the mechanistic role for EPDCs in ACM has been understudied^14^. A recent report generated Wt1-specific haploinsufficient DSP knockout mice and lineage traced EPDCs with Cre-inducible *eGFP*. scRNA-seq revealed dysregulation of epithelial-mesenchymal transition (EMT) and increased TGFβ1 signaling.^29^ Another report revealed the activation of TFAP2-dependent epithelial-to-mesenchymal transition upon *Pkp2* deletion in human-induced pluripotent stem cell (hiPSCs)-derived epicardial cells, which stimulated the expression of ECM and adipogenic genes in vitro.^15^ Here, we utilized a novel compound transgenic mouse line to delete *Pkp2* in both the epicardium and cardiomyocytes, providing the first in vivo assessment of both epicardial and cardiomyocyte involvement in *Pkp2*-deficient ACM pathogenesis. EMT markers are significantly enriched in epicardium-derived fibroblasts and pericytes, supporting a role for TFAP2A in control of fibrosis in ACM. Interestingly, although mesenchymal markers like *Vim, Tagln, Fn1*, and *Mmps* are highly elevated in Pkp2-ceKO EPDCs, Pkp2-cKO displayed more robust expression of transcriptional activators of the EMT program like *Snai1*, *Snai2*, and *Twist1,* suggesting that Pkp2-ceKO mice may exhibit more advanced disease presentation.

We detect high levels of pro-inflammatory cytokines such as *Ccl2, Ccl5, Ptx3*, and *Thbs1* in Pkp2-ceKO mice. Likewise, *Dsg2* mutant mice are reported to exhibit increased *Ccl2/Ccr2, Ccl3/Ccr5*, and *Cxcl5/Cxcr2* mRNA expression and that there is an elevation of macrophages and T cells in the fibrotic scars.^30^ These data support the recent concept that ACM pathogenesis may resemble aspects of myocarditis, which are both amplified by NF-kB signaling.^28,31,32^ Indeed, a recent report identified Ptx3 as a biomarker of myocarditis and inflammatory cardiomyopathy.^33^ It is important to note the presence of Ccl2+ and Thbs1+ fibroblasts are also found in other forms of heart disease including both ischemic and non-ischemic cardiomyopathy.^34^ The presence of Ccl2+, Thbs1+ cells has been noted in a left ventricle pressure-overload, and deleting the NF-kB activator IKKbeta in these pro-inflammatory fibroblast population attenuated Ly6C^hi^ monocyte recruitment and reduced cardiac dysfunction.^35^ Moreover, the normal human heart contains Ptx3+, Osmr+, and Il6st+ fibroblasts that also express *Ccl2, Ccl5, Thbs1, Thbs2, Mt1 and Mt2, which* may recruit c1qa+ activated macrophages.^36^ Given NF-kB is important in ACM pathogenesis, minimizing the amplification of inflammatory drive by fibroblasts may ameliorate ACM progression.

We found that, at 21 dpi, the Pkp2-deficient RV has higher recruited macrophage burden than the LV but expresses lower levels of inflammatory cytokines. This suggests a fulminant fibro-inflammatory response has developed in the RV, but is just being initiated in the LV. Ccr2+ macrophages are known to amplify pro-inflammatory macrophage recruitment and increase cardiac damage.^37^ Moreover, Ccr2+, Spp1+ and Trem2+ macrophages induce atrial and ventricular arrhythmias in mice.^38,39^ In contrast, CCR2-macrophages reduce myocardial remodeling, demonstrating the opposing role of resident and recruited macrophages in myocardial stress and damage.^24,40,41^ This temporal delay in inflammatory processes in RV vs LV introduces an interesting aspect of ACM pathogenesis that may shed light on potential therapeutic strategies to minimize the biventricular damage.

In our study, we identified increased expression of SASP factors such as *Mt1, Mt2, Tmem176b,* and *Apoe*, and various pro-inflammatory cytokines in EPDCs. SASP induces senescence in neighboring cells via paracrine signaling,^42–44^ and pro-inflammatory fibroblasts can induce senescence in other cells of the heart.^45^ Here, we find SASP is induced in various non-myocytes, which may explain the widespread SA-βgal stain in the RV of Pkp2-cKO. In addition to SASP and SA-βgal, various criteria define cellular senescence, including elevated expression of cell cycle inhibitors like Cdkn1a and Cdkn2a, nuclear lamina damage, DNA damage, chromatin abnormalities, ribosomal stress, oxidative damage, and lysosomal alterations.^46^ *Pkp2* deficient cardiomyocytes exhibit significant nuclear deformation and DNA damage along with oxidative damage.^47^ Previous reports highlight activation of the p53 pathway and increased DNA damage in ACM.^48^ In our study, we confirm the increased expression of *Cdkn1a* in both Pkp2-ceKO and Pkp2-cKO hearts, further supporting the accumulation of senescence in ACM. It is important to note that senescent populations in the heart contribute to multiple pathologies.^45,49^ The long-term effects of fibroblast senescence may be detrimental resulting in pathological fibrosis from fibroblast senescence.^50–52^ In addition, senescent cardiomyocytes contribute to pathological remodeling and dysfunction after MI, and *Cdkn2a* knockdown in myocytes prevents SASP and cardiac dysfunction.^53^ Interestingly, senescent myofibroblasts are reported to couple with cardiomyocytes via Cx43 to prolong the action potential duration and cause conduction block, which increases risk of conduction failure and arrhythmogenicity in old mice with MI.^54^ These results emphasize the various roles cellular senescence may play in cardiac pathophysiology, potentially including arrhythmogenesis.

In our scRNA-seq, we observe an over-abundance of B cells in Pkp2-ceKO mice. Although B cell depletion did not yield a difference in cardiac physiology, it did ameliorate LV fibrosis at 21 dpi. This result suggests B cell depletion may modulate inflammatory and fibrotic cascades in the early damage response. B cells are known to be mostly of peripheral origin, but some studies report subsets of B lymphocytes that reside within the heart.^55, 56^ It has also been noted that B cells may play a protective role in reducing inflammation and fibrosis from acute cardiac insult.^56^ However, a previous study analyzed B cell interactions with other immune cells, and found only minor dysregulation of inflammatory pathways in myocardial B cells during ACM.^57^ Taken together, these data may explain why B cell depletion does not show a more robust impact in later stage ACM when the disease has progressed beyond the RV.

### Study Limitations

There are number of limitations of the study that should be addressed. First, the *Pkp2* KO mouse model does not capture all aspects of the ACM pathology. For example, instead of the fibrofatty deposits characteristic of human ACM, Pkp2-cKO and Pkp2-ceKO mice exhibit fibrosis without fat accumulation. Other models of ACM, such as *Dsp* KO mice capture the fibrofatty deposition in the myocardium, suggesting potential differences in the molecular pathways elicited by various ACM mutations. Another possibility is that rapid disease progression in the Pkp2-cKO mouse may obscure more subtle aspects of the disease phenotype. Second, scRNA-seq data was obtained from only male mice given the male predominance of disease and constraints on resources. However, validation studies were performed in female mice suggesting our observations are generalizable between sexes. Third, we did not perform scRNA-seq on cardiomyocytes, which prevented a direct comparison of communication between cardiomyocytes and non-myocytes by CellChat. However, integrating our previous bulk RNA-seq data from whole ventricle tissue to our current scRNA-seq suggests myocyte contributions either occur before 21 dpi or are via a mechanism that is independent of gene dysregulation; potentially via DAMP production.

## Conclusions

Our study reveals a role for epicardium-derived fibroblasts in Pkp2 deficient ACM. Pkp2 deletion leads to an accumulation of senescent and pro-inflammatory fibroblasts, which correlates with the recruitment and activation of immune cells in the heart. This study also sheds light on the epicardium as an initial driver of inflammation in ACM and suggests a temporal window when anti-inflammatory interventions may ameliorate LV remodeling.

## Nonstandard Abbreviations and Acronyms

ACM: Arrhythmogenic Cardiomyopathy
CM: Cardiomyocyte
CVD: Cardiovascular Disease
Dpi: Days post-tamoxifen injection
EPDCs: Epicardial Progenitor Derived Cells
DAMP: Damage-associated molecular pattern
DSC2: Desmocollin-2
DSG2: Desmoglein-2
DSP: Desmoplakin
GFP: Green Fluorescent Protein
JUP: Plakoglobin
LV: Left Ventricle
PKP2: Plakophilin-2
PKP2-cKO: Cardiomyocyte-specific knockdown of Plakophilin-2
PKP2-ceKO: Cardiomyocyte and Epicardium-specific knockdown of Plakophilin-2
PKP2-eKO: Epicardium-specific knockdown of Plakophilin-2
RV: Right Ventricle
SASP: Senescence-Associated Secretory Phenotype
SCD: Sudden Cardiac Death
scRNA-seq: Single-Cell RNA Sequencing

## Acknowledgement

We thank the University of Rochester Medical Center Flow Cytometry Core for assistance with cardiac immune cell quantification, and the Genomics Research Center for their assistance with scRNA-seq.

## Supplemental Material

Supplemental Methods Supplemental Table 1

Supplemental Figures 1-7

## Novelty and Significance

### What is known?

- Pkp2 deficient arrhythmogenic cardiomyopathy (ACM) is associated with myocardial inflammation and the accumulation of fibro-fatty tissue.
- Disease progresses from the epicardium-to-endocardium, suggesting epicardial cells may contribute to tissue remodeling.

### What new information does this article contribute?

- Using single cell transcriptomics, we defined how non-myocyte phenotypes change in mice lacking Pkp2 in cardiomyocytes and epicardium-derived cells, compared to controls.
- Pkp2 deficient epicardium-derived fibroblasts become senescent and acquire the senescence-associated secretory phenotype (SASP), characterized by an abundance of Ccl2, Ccl5, Thbs1, Ptx3, and Il6.
- Accumulation of pro-inflammatory fibroblasts corresponds to a significant immune cell burden in Pkp2 deficient mice, characterized by the recruitment and activation of macrophages, B cells and T cells.
- B cell delays the initiation of fibrotic and inflammatory responses but does not significantly improve end stage heart function.

Our findings present a novel mechanism in the pathogenesis of Pkp2-deficient Arrhythmogenic Cardiomyopathy (ACM). We found that Pkp2-deficient epicardium-derived cells are a source of cytokines that may increase the inflammatory burden in ACM. Pkp2 deficient epicardium-derived fibroblasts become senescent and acquire the senescence-associated secretory phenotype (SASP), characterized by an abundance of Ccl2, Ccl5, Thbs1, Ptx3, and Il6. The epicardium-directed pro-inflammatory program follows a right ventricle to left ventricle predominance as disease progresses and correlates with an abundance of macrophages, T-cells and B-cells. While end stage disease is not affected by B cell depletion, the fibrotic and inflammatory responses are significantly delayed. Through better understanding the timing and immunological basis of ACM pathogenesis, in conjunction with the importance of epicardial contributions to inflammation, this study may help to accelerate ACM treatment strategies that target the immune system.

