## Supplemental Methods and Figures for "Epicardial contributions to fibro-inflammatory signaling in a Pkp2-deficient arrhythmogenic cardiomyopathy model"

**Animal Models**

All experiments using animals were approved by the University Committee on Animal Resources at the University of Rochester. Experiments involved using male and female mice with the following genotype: wildtype controls (Myh6$^{MerCreMer}$;Wt1$^{CreERT2}$;Pkp2$^{+/+}$), PKP2-cKO (Myh6$^{MerCreMer}$; PKP2$^{fl/fl}$), and PKP2-ceKO (Wt1$^{CreERT2}$; Myh6$^{MerCreMer}$;Pkp2$^{fl/fl}$). Myh6$^{MerCreMer}$ mouse lines was obtained from (JAX #005657).[18] Wt1$^{CreERT2}$ mice were used to efficiently label the epicardium and its derivatives as previously described and were obtained from (JAX #010912).[11,17,20] PKP2$^{fl/fl}$ line was obtained from Dr. Mario Delmar.[16] For lineage tracing experiments, mice were bred to the Rosa26$^{mTmG}$ line, which labels cells via Cre-mediated recombination. *Rosa*$^{mTmG/mTmG}$ mouse strain was purchased from (JAX #007576). Mice were genotyped when they were pups and regenotyped after each experiment to confirm correct genotypes. The list of primers for each specific gene is in Supplemental Table 1. Mice were injected at 8-12 weeks of age with 100mg/kg of tamoxifen (Sigma-Aldrich, St. Louis, MO) for 4 days to induce the Cre-mediated knockdown.

**Echocardiography**

A baseline echocardiographic analysis was performed before Tamoxifen injection, with subsequent echoes at 21- and 42-days post-Tamoxifen injection (dpi) in all genotypes. Mouse were anesthetized with isoflurane before the echocardiographic measurement. Echocardiographic measurements were performed using a Vevo3100 echocardiography machine (VisualSonics, Toronto, Canada) and a linear-array 40MHz 2 transducers (MS-550D). LV systolic and diastolic measurements were captured in M-mode from the parasternal short axis. RV systolic and diastolic measurements were captured in M-mode from the parasternal long axis. All echocardiography was performed by an investigator who was blinded to different genotypes.

**Picrosirius Red staining and fibrosis measurement**

After 21 or 42 dpi, mice were anesthetized by injecting 0.5 mL of 13mg/mL of ketamine and 0.88 mg/mL of xylazine solution. After confirmation of full unconsciousness, the chest cavity was opened and the heart was perfused with calcium-and magnesium-free phosphate-buffered saline (PBS), followed by perfusion with 10% neutral buffered formalin solution and fixation overnight. After fixation, the hearts were processed, embedded in paraffin, and were sectioned into 5 um sections. For staining, slides were deparaffinized in a series of xylenes, followed by 3-minute incubations in 100% ethanol (EtOH, 3x), 95% EtOH (1x), and then placed in distilled water. The Picrosirius red stain (Abcam, Cambridge, MA) was used to stain for collagen deposition in the heart according to the manufacturer’s instructions. Stained slides were imaged by bright field imaging using an Olympus BX51 at 1.25x magnification. To evaluate fibrosis, total areas of each RV and LV ventricles were defined in FIJI ImageJ software. The fibrosis was quantified by calculating the ratio of the total area of the positive red pixels in each ventricle to the total area of each ventricle.

**Epicardial-Endocardial fibrosis gradient measurement**

For analysis of epicardial to endocardial gradient of fibrosis, QuPath 0.5.1 was used to read Picrosirius red stained heart sections. Two separate AI-trained segmentation models were created for the fibrosis and the background to allow for independent retraining of the models. These segmentation models were trained over multiple images and validated with images not used for training. The background segmentation consists of an inside and an outside annotation that has a nearly identical border, but allow the annotations to be separated into one that outlines the inside chambers of the heart wall, and another that only outlines the outside of the heart wall. Separate annotations are needed to calculate the distance from the fibrosis to both the inside and the outside annotations and to measure the approximate width of the heart wall. Downsampling the annotations helped create inside annotations with closed loops over both chambers of the heart. The annotations are split by removing holes of a specific size and then removing the intersection of the two annotations. Before applying either AI segmentation model, the user can manually annotate the fibrotic region of the heart and then the AI segmentation will be restricted to the annotated area. The image is segmented into the three main parts, fibrosis, an inside annotation, and an outside annotation. To facilitate heart width measurements at multiple fibrotic regions, the fibrosis segmentation is then clustered into superpixels. For every superpixel, the signed distance to both the inner and outer annotations is calculated. Signed distances allow for distinction between fibrosis located inside or outside the heart. Fibrosis superpixels located outside of the heart can be ignored. The approximate width of the heart can be calculated from the sum of the absolute value of the two distances.

Picrosirius red stain of RV LV from Pkp2-cKO hearts (n=5) were used for the analysis. All the pixels counts were collected and reported in histogram to summarize the gradient.

**Langendorf Isolation of non-cardiomyocytes**

After anesthetizing the mice, the chest was opened to reveal the aortic arch. A small incision was made in the upper portion of aortic arch, and a 22-gauge cannula (Radnoti LLC, Covina, CA) was inserted and sutured to hold it in place. After aortic cannulation, the heart was perfused with Perfusion buffer at 37°C, which consists of 120 mM of NaCl, 14.8 mM of KCl, 0.6 mM of $KH_2PO_4$, 0.7 mM of $Na_2HPO_4$, 2.5 mM $MgCl_2$, and 10mM HEPES with pH adjusted to 7.35 with NaOH. Then, the heart was perfused with Digestion buffer at 37°C, which consists of 15,000 units of collagenase type II (Worthington Biochemical Corporation, Lakewood, NJ) dissolved in Perfusion Buffer, on EZ Myocyte/Langendorff Isolated Heart System (Radnoti LLC, Covina, CA) for 6-8 minutes. Upon completion of digestion, the atria and great vessels were removed and only the ventricles were isolated. Then, it was placed in a petri-dish with Digestion buffer and forceps were used to gently shred the ventricles into small pieces. Stop buffer, which consists of 10% FBS and 12.5 mM $CaCl_2$, was added and the ventricles were pipetted up and down 20 times with plastic pipette. Next, cardiomyocytes were allowed to settle to the bottom of a 15mL conical tube by gravity after 15 minutes of incubation in the 37°C water bath, with gentle tapping every 5 minutes. Supernatants with non-myocytes were collected with pipetting and the remaining cardiomyocytes were discarded. Non-myocyte fractions were sent to the Genomic Research Core (GRC) at the University of Rochester Medical Center (URMC) to generate the single cell libraries on the day of isolation.

**Single cell library preparation and processing**

scRNA-sequencing data has been deposited in the Gene Expression Omnibus (GEO) database under accession code GSE300141. All the scRNA-seq library preparation, processing, and analysis were conducted in the Genomic Research core (GRC) at URMC. Single cell libraries were generated from non-myocytes acquired by differential adherence to plastic following Langendorff digestion of mice of the following genotype (n=2): wildtype Cre controls (Myh6$^{MerCreMer}$;Wt1$^{CreERT2}$;Pkp2$^{+/+}$), PKP2-cKO (Myh6$^{MerCreMer}$; PKP2$^{fl/fl}$), and PKP2-ceKO (Wt1$^{CreERT2}$;Pkp2$^{fl/fl}$). Prior to capture using the 10x Genomics Chromium controller (10x Genomics), the number of cells was quantitated (TC20 Automated Cell Counter, Bio-Rad) and cell viability was assessed via the trypan blue exclusion test of cell viability. Only cell populations exhibiting greater than 80% viability were used. All cells were loaded to maximize the number of single cells acquired using the Chromium single Cell 3’ Reagent Kit. Libraries were prepared according to manufacturer’s instructions using the Chromium Single Cell 3’ Library and NextGEM v3.1 bead kit (10x Genomics). CellRanger v3.1.0 was used to demultiplex each capture, process base-call files to fastq format, and perform 3’ gene counting for each individual cell barcode with mouse reference data set incorporating custom tdTomato and EGFP sequences (mm10, v 3.0.0).

**Cell filtering and cell-type annotation and clustering analysis.**

Seurat v4.1.0 within R v4.1.1 was used to integrate and process the samples. Samples were imported, keeping cells only with a minimum of 200 unique genes detected and genes detected across at least 3 cells. Cells were additionally filtered for cells where the total captured content was less than 15% mitochondrial. Samples were merged using "merge", normalization performed regressing out mitochondrial content, and clustering performed at a resolution of 0.3 and 0.8. Cell typing was performed via visualization of expression for canonical markers. Seurat was used for the visualization of markers, specifically the VlnPlot and FeaturePlot functions. Transcription factor analysis was performed using EnrichR v3.0 against the ChEA_2022 database for markers determined via the FindAllMarkers function.

**Developmental trajectory and prediction of cell-fate determinants.**

Epicardial populations were identified, isolated, and reclustered, keeping PKP-cKO cells that were $eGFP^-$ and Cre control & PKP2-ceKO cells that were $eGFP^+$. Epicardial populations were clustered to a resolution of 0.6. The Seurat object was converted to a SingleCellExperiment using SingleCellExperiment v1.14.1 for use in trajectory analysis. Slingshot v2.2.1 was used for analysis, selecting the cluster overrepresented for PKP2-ceKO as the terminal endpoint (via the end.clus parameter). From this, four lineages were generated for further analysis. A negative binomial general additive model was fitted to the lineages using tradeSeq v1.8.0 (with nknots = 7) to allow for statistical inference across the trajectory. associationTest was used to perform differential expression with “lineages = T” and “contrastType = ‘end’”. Lineages were separately analyzed with p-value adjustment performed using R’s built in p.adjust function. Genes from each lineage with a meanLogFC greater then 1 were used for pathway enrichment against the GO_Biological_Process_2023, KEGG_2019_Mouse, and WikiPathways_2024_Mouse libraries via EnrichR v3.0.

##### Processing and Integration of PKP2 RNA-Seq

Quality filtering and adapter removal was performed using FastP v.0.23.1. Processed reads were then mapped to the mouse reference genome (GRCm39 + gencode M31) (https://www.gencodegenes.org/mouse/release_M31.html) using STAR_2.7.9a. Reads mapping to genes were counted using subread featurecounts v2.0.1 with “-s 2”. Differential expression analysis was performed using DESeq2-1.34.0 with an adjusted P-value threshold of 0.05 within R v4.1.1. Gene ontology analyses were performed using the EnrichR-3.0 package. The bulk RNA-Seq dataset was integrated into the single cell RNA-Seq dataset by importing the raw counts as a Seurat object. From there, the datasets were merged using the merge function, normalized using SCTransform, PCA run, and UMAP run using the first 20 dimensions of PCA. The Seurat object was then saved for use within cell-cell communication analysis.

##### CellChat for cell-cell communication analysis

The dataset was imported within R v4.3.1 using Seurat v5.0.3.9911. CellChat v2.1.2 was used for analysis. A CellChat object was created from the Seurat object using createCellChat. Mouse was chosen as the CellChat reference using CellChatDB.mouse. The following functions were performed iteratively on the object to determine ligand and receptor interactions within the cell populations: subsetData, identifyOverExpressedGenes, identifyOverExpressedInteractions, computeCommunProb (using type = “triMean”), computeCommunProbPathway, aggregateNet, and netAnalysis_computeCentrality(using slot.name = “netP”). The netVisual_bubble and netVisual_chord_gene functions were used for visualization.

**Cytokine Array**

At 21 days dpi, mice were anesthetized with isofluorane and the blood was collected by retro-orbital bleeding using heparinized capillary tubes (Fisher Scientific, Waltham, MA) and placing blood in collection tubes (Fisher Scientific, Waltham, MA) and was allowed to coagulation for 1 hour before centrifugation of samples for 15 minutes at 2000g. Plasma was collected after centrifugation The top layer or serum was removed from each sample and placed at -80ºC until ready to use. A total of 120µL of serum was used by pooling equal volumes of sera in each Cre Control (n=3) and PKP2-cKO (n=4) mice. Cytokine Array was performed according to the manufacturer’s instructions (R&D Systems, Minneapolis, MN). Cytokines were detected by chemiluminescence and exposure for up to 15 minutes using the Chemidoc Imaging System (Biorad, Philadelphia, PA). Relative intensity of signal between WT and KO was analyzed with FIJI ImageJ Software (NIH, Bethesda, MD).

**Flow Cytometry**

Hearts of male PKP2-ceKO mice at 10 dpi (n =5) and 21 dpi (n =4) were processed to single cell suspensions following Langendorff digestion (as described above). CM were removed by gravity sedimentation, and supernatant containing non-CM was filtered through 50mm cell strainer and pelleted at 250g for 3 minutes at 4°C. Red blood cells were lysed with ACK lysis buffer for 5 minutes at room temperature and cells were again pelleted as above, followed by resuspension in 1.5 mL of FACS buffer (PBS + 2% Fetal calf serum + 2mM EDTA). For B cell quantification, a panel of antibodies against CD45-PerCP-Cy5.5 (BioLegend, Cat#103131), and CD19-PE (BioLegend, Cat#152407), were incubated with cells on ice for 30 minutes at 1:1500 dilution each. Cells were then pelleted as above and resuspended in 1.2mL FACS buffer and sorted on a BD FACS Aria II in the Flow Core at the University of Rochester Medical Center.

**B cell Depletion**

B-cell depletion followed the protocol conducted previously[55]. anti-CD20 antibody (BioLegend, Cat #152104) or IgG2b isotype control (BioLegend, Cat #400671) was injected retro-orbitally (100μg) a day prior to first dose of tamoxifen (~ 8-12 weeks of age) in PKP2-ceKO mice (n=4 mice per group). On 7 and 21 dpi, blood was collected from tail bleeding and flow cytometry was conducted to confirm a successful B cell depletion. Heart was isolated at 21 days dpi and was used for qPCR or histology. Echocardiography was conducted as described previously at 21 dpi in a separate larger cohort (IgG, n=10; CD20, n=11). Picrosirius red stains were done in each treatment group as described previously (IgG, n=6; CD20, n=7). The remainder of the heart was used for qPCR analysis (IgG, n=3; CD20, n=3).

**RNA isolation, cDNA synthesis, and RT-qPCR**

After perfusion of the hearts with PBS, the right ventricle free wall was separated by using the scissors to cut along the border where the septum lies. Septum was kept along with the left ventricle. The ventricles were each placed in the Eppendorf tubes and were immediately frozen by placing them in liquid nitrogen. Ventricles were kept in -80°C fridge for long-term storage.

For RNA isolation, frozen ventricles were weighed and hammered to fine power using for further processing. Powdered hearts were added with TRIzolTM Reagent (Thermo Fisher Scientific) and was homogenized with Pro Scientific PRO200 Homogenizer to ensure maximal yield of RNA. Then, the RNA was isolated according to the manufacturer's instructions. RNA was treated with the TURBO DNA-free™ Kit (Thermo Fisher Scientific) to eliminate genomic DNA. Purified RNA amounts were normalized across samples and then were made into cDNA using iScript cDNA Synthesis Kit (Biorad). RT-qPCR was performed with cDNA, primers and IQ SYBR Green Supermix (Biorad) in Biorad CFX Connect and CFX Duet Real-Time qPCR system (Biorad). Data was analyzed using the ΔΔC(t) method. The list of primers for each specific gene is in Supplemental Table 1.

**SA-βgal Staining**

After perfusion of the hearts with PBS, the hearts were embedded in the OCT (Sakura) and were frozen on a metal stand submerged in isopentane (Thermo Fisher) cooled with liquid nitrogen. Then, the frozen tissue blocks were kept in -20°C fridge and the slides were sectioned the day after in 5 um thickness with Leica CM 1950 Cryostat. The slides were kept for long-term storage at -80°C. PKP2-cKO and Myh6-Cre+ control mice were used with n=3/genotype. SA-βgal staining was done according to the manufacturer's instructions (Sigma). Frozen slides were slowly thawed at 4°C for 1 hour to minimize damage in integrity from rapid thawing. X-gal solution was warmed at 37°C for 1 hour to minimize the formation of crystal aggregates. Slides were washed for 5 minutes in PBS three times to wash out the residual OCT. After lightly fixing the slides in the fixation buffer for 7 minutes, slides were washed for 5 minutes in PBS three times. Then, slides were incubated in the X-gal solution for 48 hours in 37°C incubator without $CO_2$ and protected from light. After 48 hours, slides were washed for 5 minutes in PBS three times. Stained slides were imaged by brightfield imaging using an Olympus BX51 at 20x magnification.

**Statistical Analysis**

Replicates in this study were biological (not technical) unless specifically noted otherwise, and all data are reported as mean +/- SD. Statistical analyses were performed using Prism 9.1.1. (GraphPad). For comparison between two groups that were not normally distributed, non-parametric unpaired, two-tailed Mann Whiteny test was conducted. If normally distributed, the comparison was performed using unpaired, two-tailed t-tests with Welch's correction for unequal standard deviations. To compare the differences among three or more groups of data that are not normally distributed, we used the Kruskal-Wallis test with Dunn's post-hoc analysis. For groups of data that were normally distributed, one-way ANOVA was used with Holm- Šidák post-hoc analysis.

Geisser-Greenhouse correction was used for repeated measures two-way ANOVA. Post-hoc comparisons were made using the Tukey test for two-way repeated measures ANOVA where three or more groups were compared. $p < 0.05$ was considered significant.

**Supplemental Table 1.** List of genotyping and qPCR primers

| Gene Name | Forward Primer (5' -> 3') | Reverse Primer (3' -> 5') |
|---|---|---|
| **Genotyping** | | |
| Pkp2 | GAG AGA TTA CTG GCA TAC ACT ACT GG | CTG CCG TCC AAC AAA GTC ATT CTC |
| Wt1-CreERT2 | ATC GCA GGA GCG GAG AAC | GAA GGG TCC GTA GCG ACA (Wildtype)<br>GCA AAC GGA CAG AAG CAT TT (Transgenic) |
| Myh6-MerCreMer | CTA GGC CAC AGA ATT GAA AGA TCT (Wildtype)<br>ATA CCG GAG ATC ATG CAA GC (Transgenic) | GTA GGT GGA AAT TCT AGC ATC ATC C (Wildtype)<br>AGG TGG ACC TGA TCA TGG AG (Transgenic) |
| Rosa-mTmG | CTC TGC TGC CTC CTG GCT TCT<br>(Wildtype)<br>TCA ATG GGC GGG GGT CGT T (Transgenic) | CGA GGC GGA TCA CAA GCA ATA<br>(Wildtype) |
| **qPCR** | | |
| Pkp2 | TGT TTT GGC AGC CGA AGT AG | AAT GCC ACA GCC ACT CAA TG |
| Cd19 | GGC ACC TAT TAT TGT CTC CG | GGG TCA GTC ATT CGC TTC |
| Postn | GCA AAC CAC TTT CAC CGA CC | CGT TGG TCC ATG CTC AGA GT |
| Col1a1 | TAG GCC ATT GTG TAT GCA GC | ACA TGT TCA GCT TTG TGG ACC |
| Col1a2 | AGC AGG TCC TTG GAA ACC TT | AAG GAG TTT CAT CTG GCC CT |

| | | |
|---|---|---|
| Col3a1 | TAG GAC TGA CCA AGG TGG CT | GGA ACC TGG TTT CTT CTC ACC |
| Ccl2 | CAC TCA CCT GCT GCT ACT CA | GAG CTT GGT GAC AAA AAC TAC AGC |
| Thbs1 | GGT AGC TGG AAA TGT GGT GCG T | GCA CCG ATG TTC TCC GTT GTG A |
| Ptx3 | CCT GCG ATC CTG CTT TGT G | GGT GGG ATG AAG TCC ATT GTC |
| Ccl7 | GTC CCT GGG AAG CTG TTA TCT TCA | GCA GAC TTC CAT GCC CTT CT |
| Cdkn1a | GTT CCT TGC CAC TTC TTA CCT | TCA TCC TAG CTG GCC TTA GA |
| Apoe | AAC AGA CCC AGC AAA TAC GCC | CTC ATT GAT TCT CCT GGG CC |
| Tmem176b | ACT CCA GCT AGA ATT GCC ACA G | CAT CAG CAT CCA CAT CCA CC |
| Adgre1 | AAG ACT GAC AAC CAG ACG GC | CTG CCT CCA CTA GCA TCC AG |
| Ccr2 | TTA CAC CTG TGG CCC TTA TTT | CTG AGT AGC AGA TGA CCA TGA C |
| Cx3cr1 | CAG CAT CGA CCG GTA CCT T | GCT GCA CTG TCC GGT TGT T |
| Spp1 | TCT CCT TGC GCC ACA GAA TG | TGT GGT CAT GGC TTT CAT TGG A |
| Trem2 | TGG GTC CCC GAG GAG TCA TC | AAA CTT GCT CAG GAG AAC GC |
| 18s | GTA ACC CGT TGA ACC CCA TT | CCA TCC AAT CGG TAG TAG CG |

### Supplemental Figure 1

A

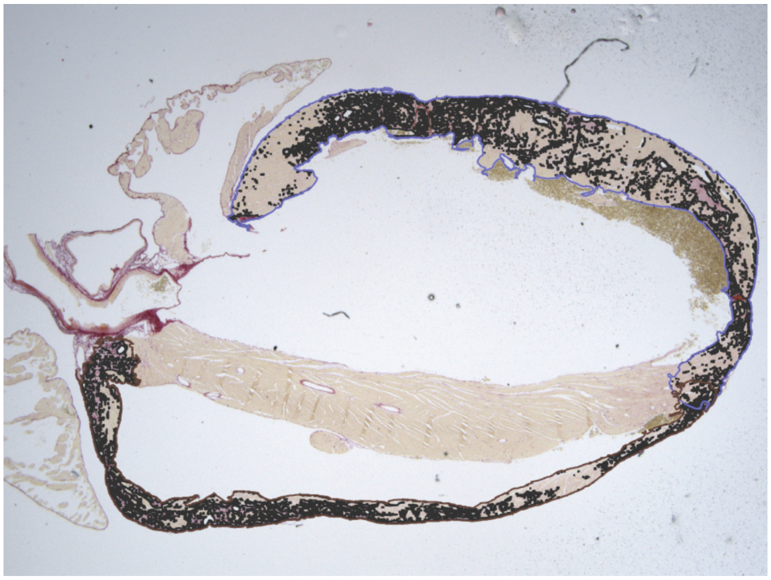

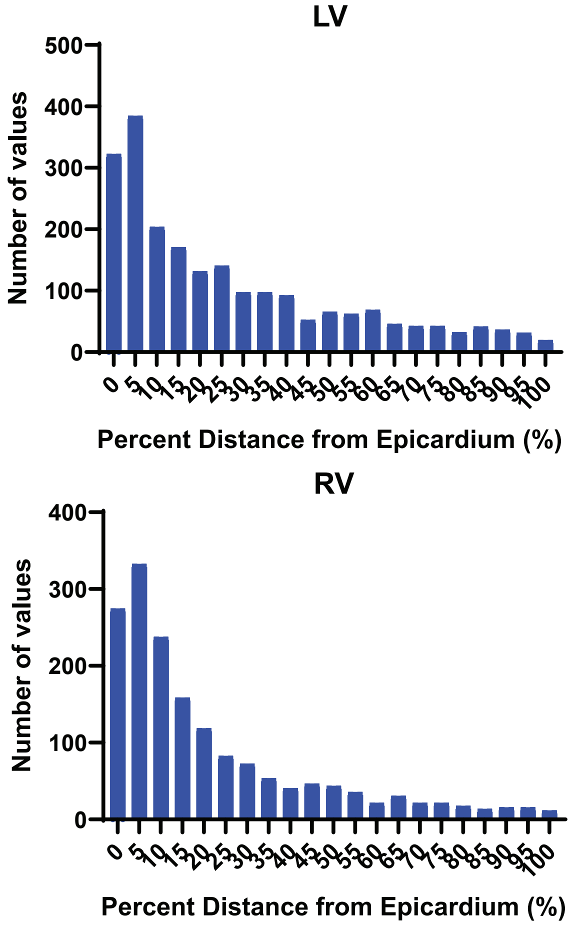


B

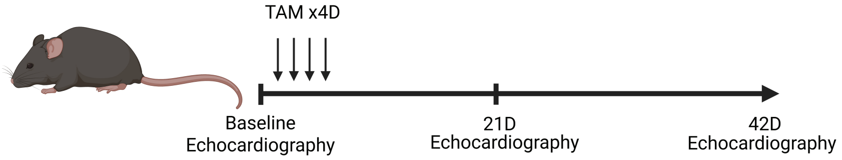


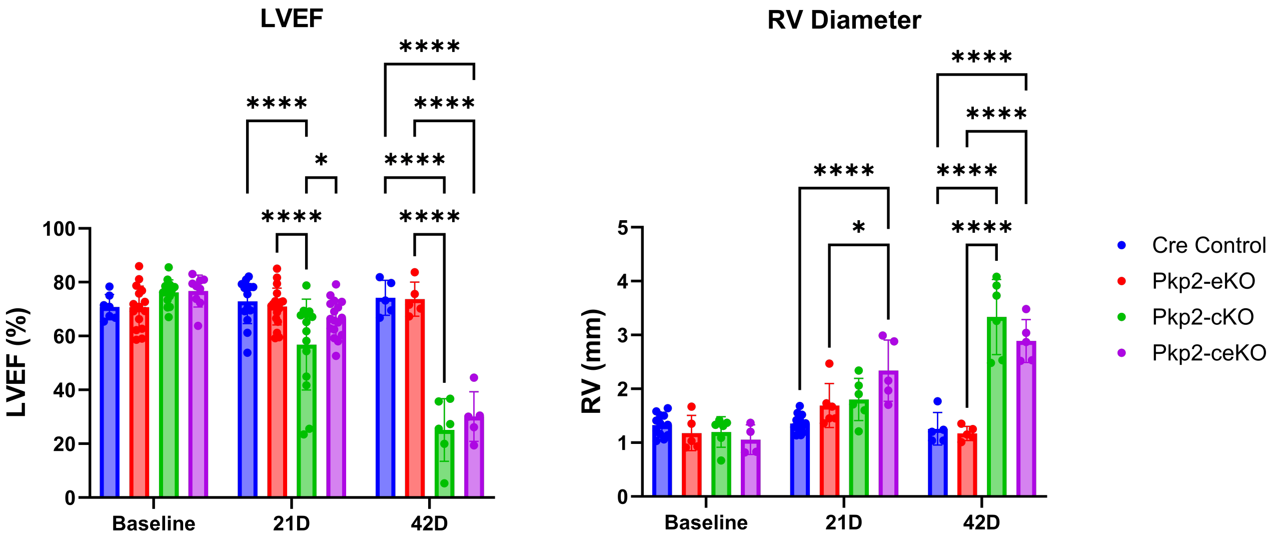

#### Supplemental Figure 2

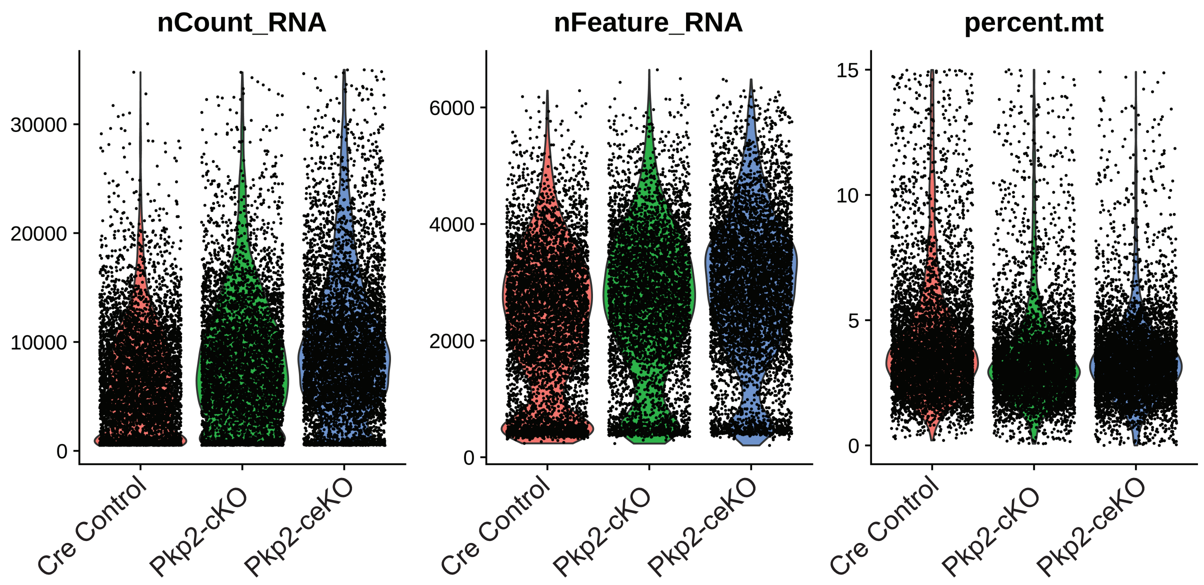

### Supplemental Figure 3

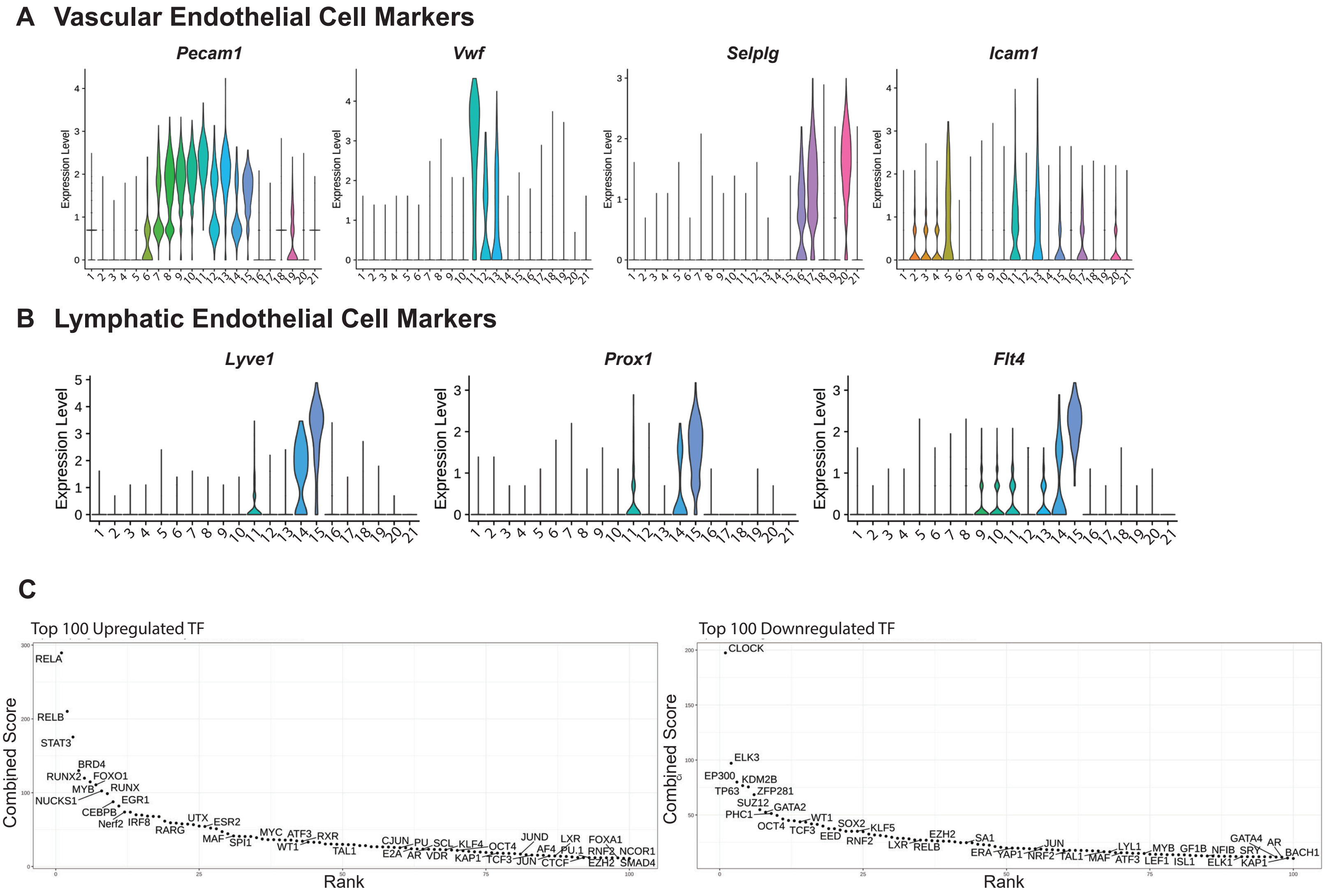

#### Supplemental Figure 4

### A

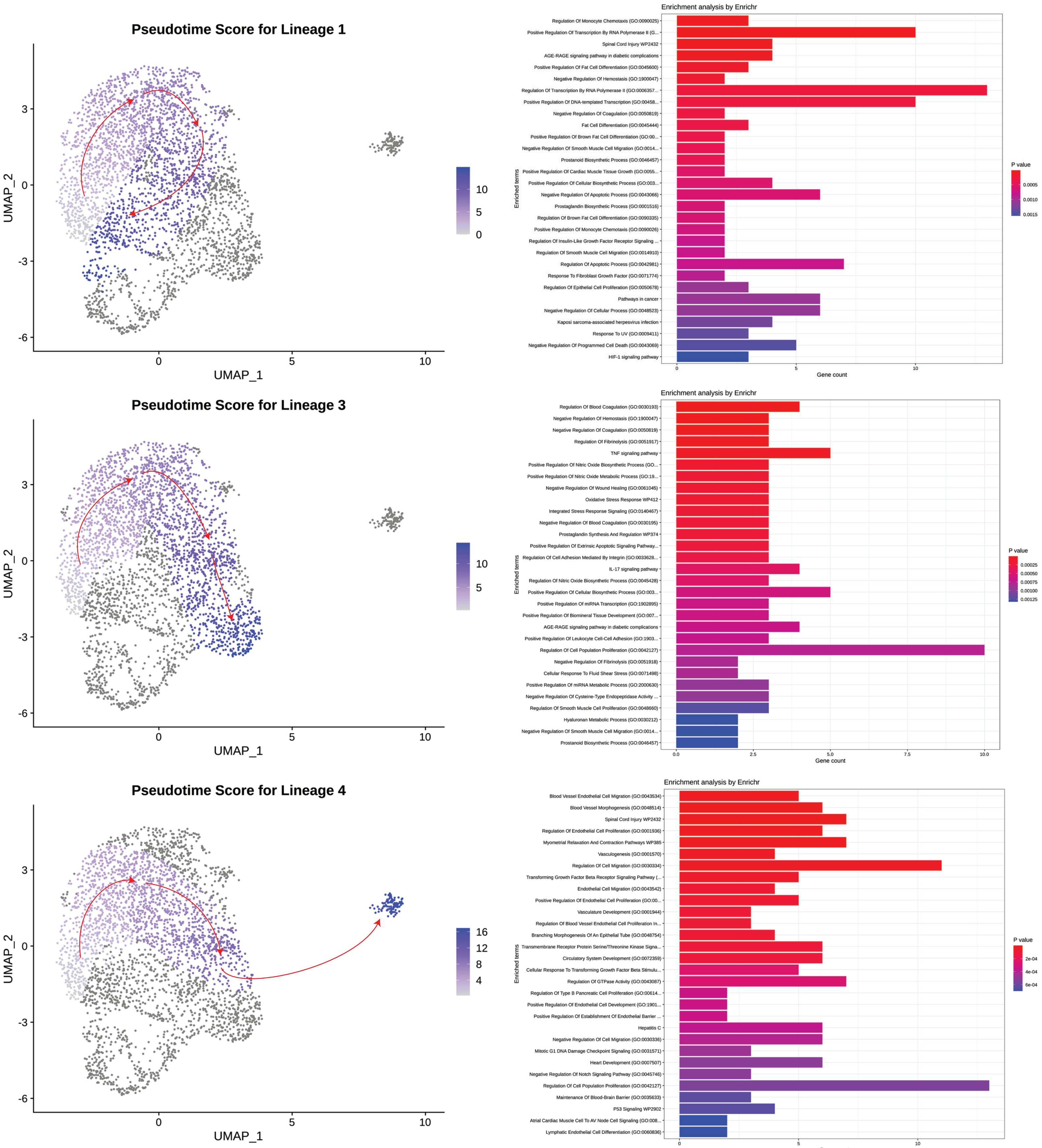

### Supplemental Figure 5

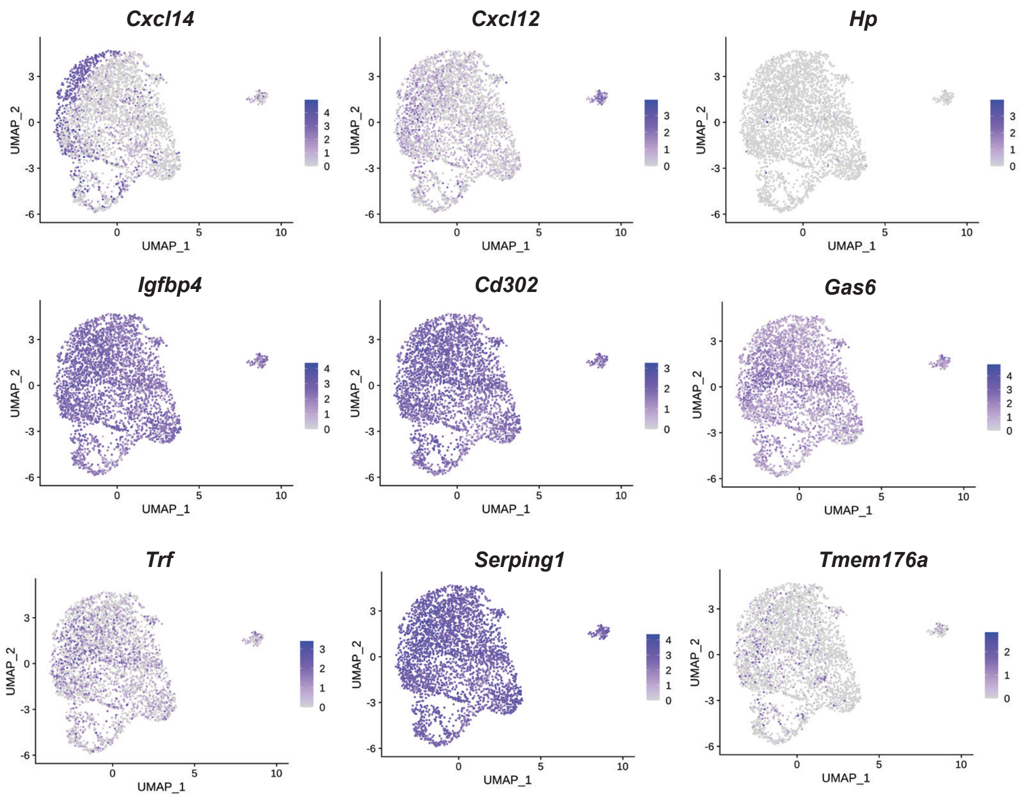

### Supplemental Figure 6

#### A Macrophage/Monocyte Markers

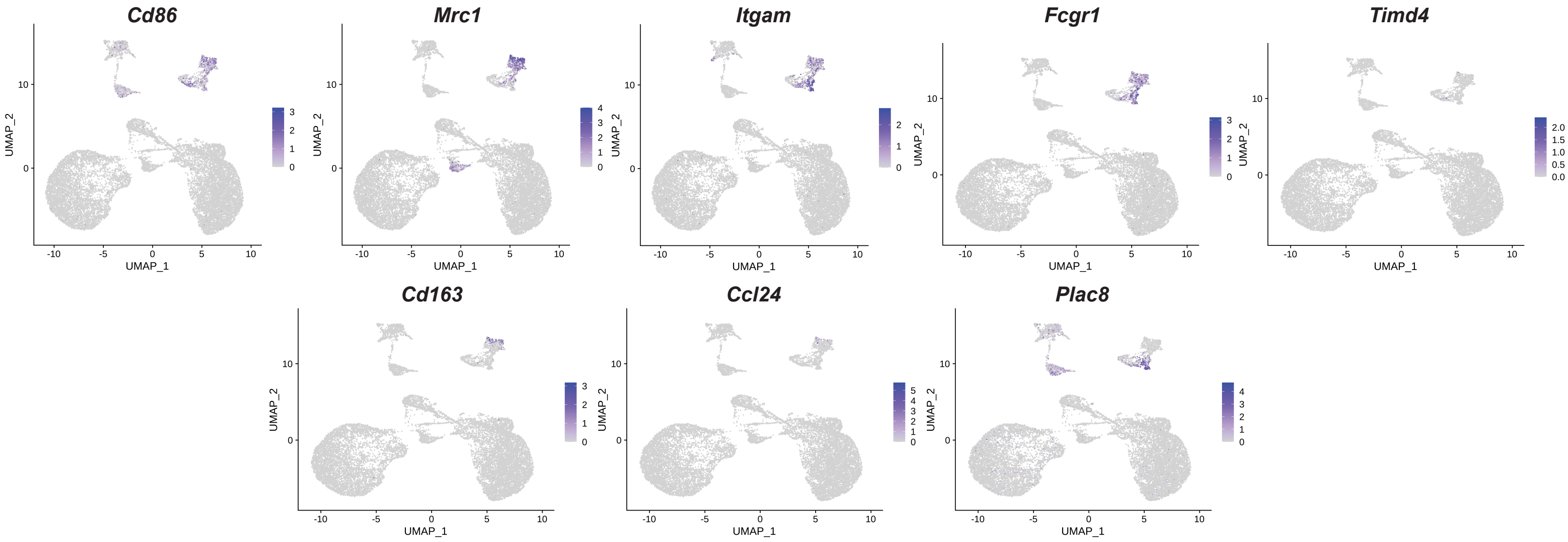


#### B B Cell Markers

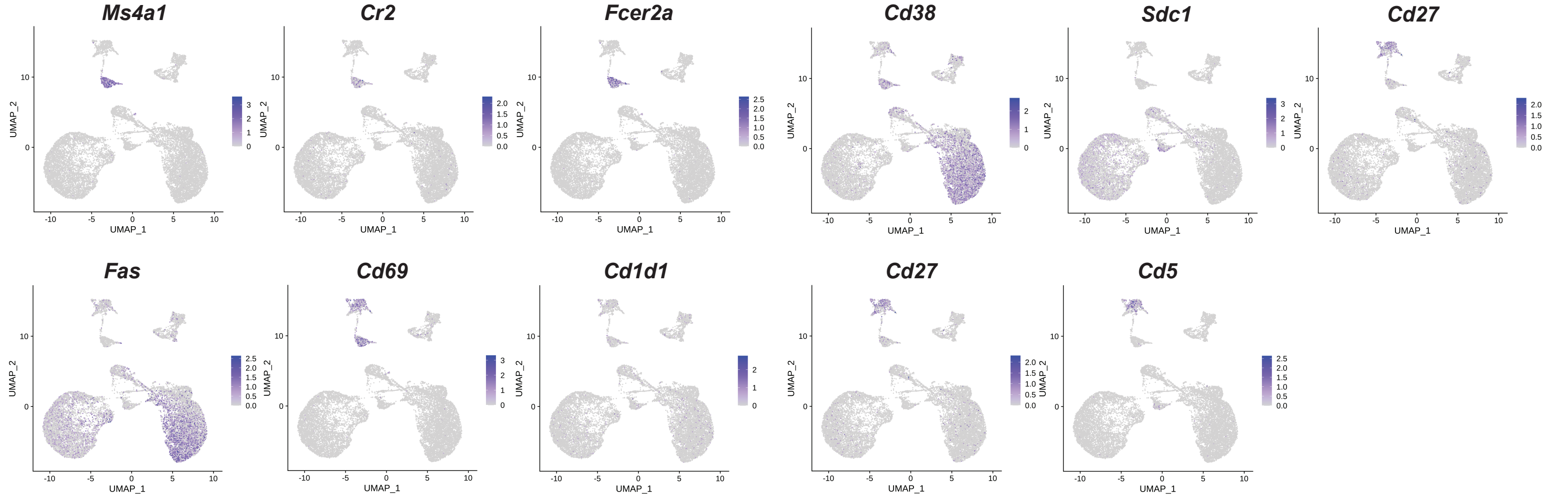


#### C T Cell Markers

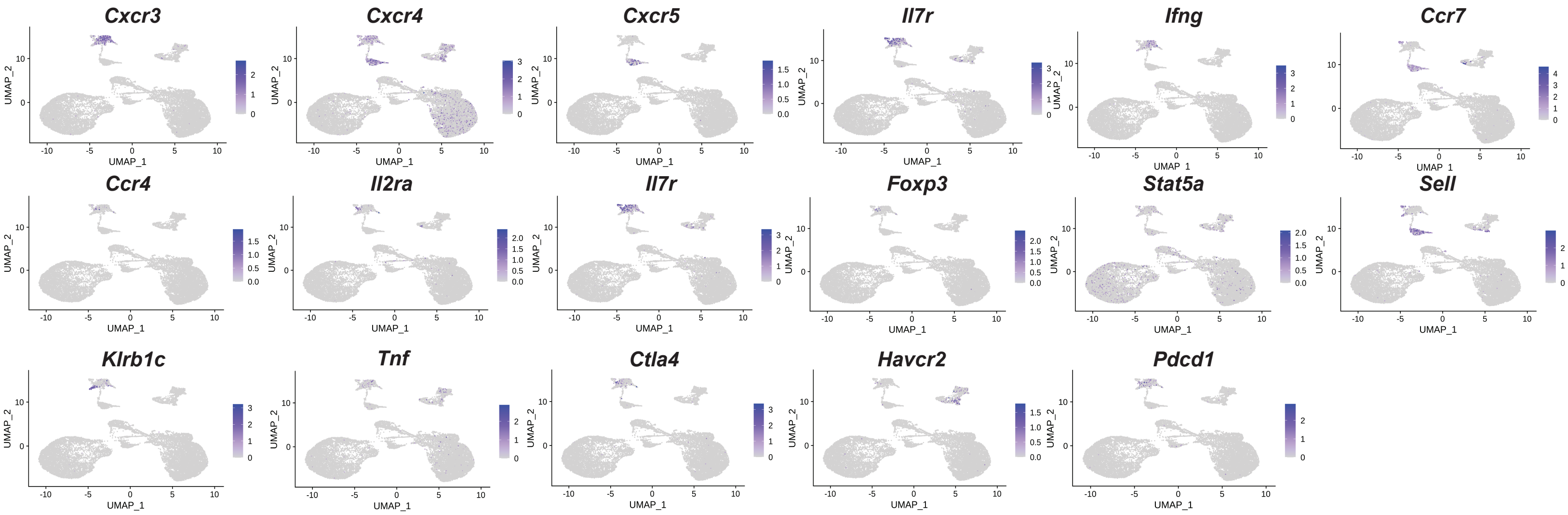

### Supplemental Figure 7

## A

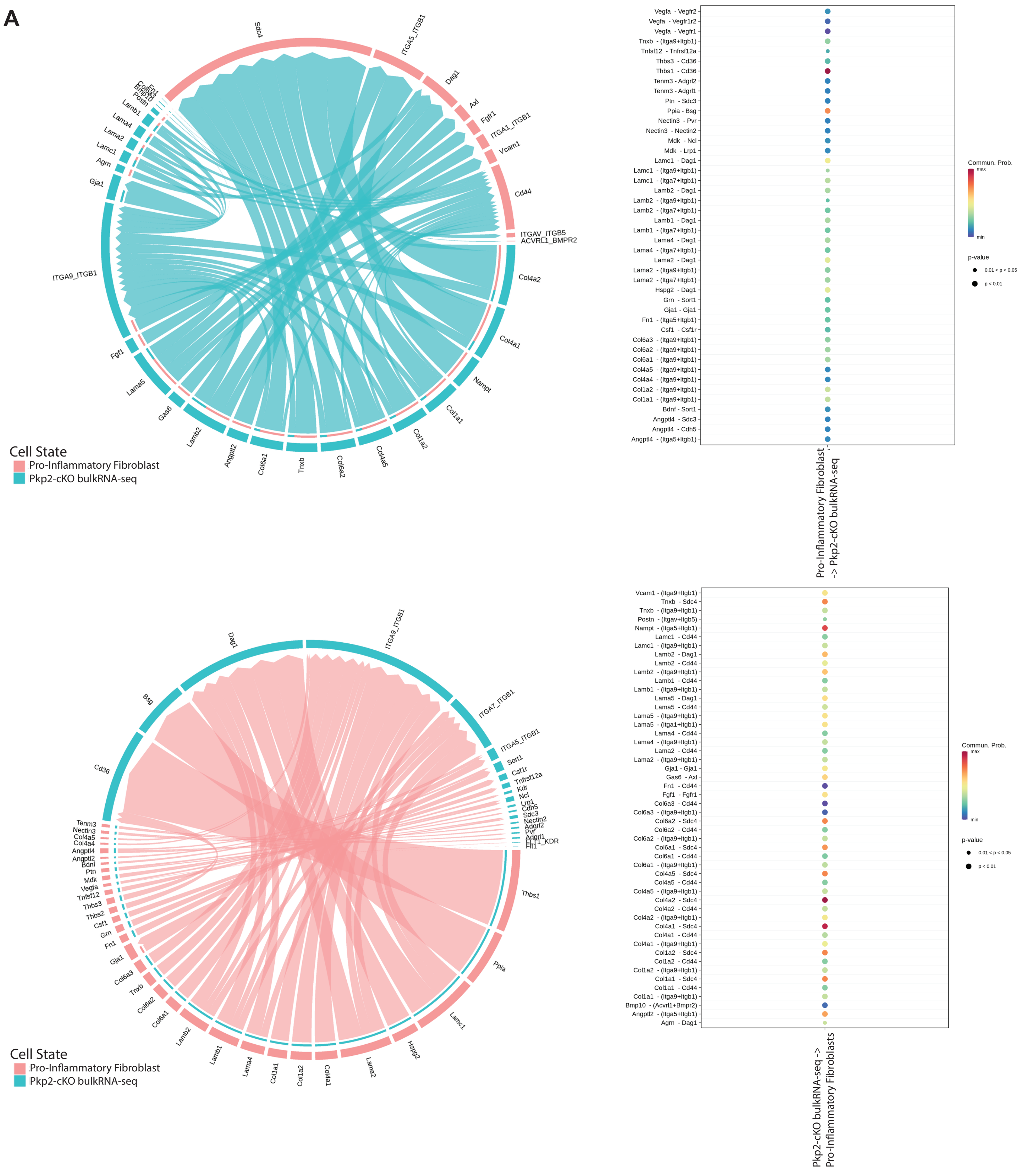

**Supplementary Figure Legends**

**Supplementary Figure 1. A)** Representative Picrosirius stained heart with red pixels defined in each ventricle and s of quantification of epicardial to endocardial gradient of fibrosis in LV and RV from Picrosirius Red stained Pkp2-cKO hearts. **B)** Schematic describing experimental timeline for tamoxifen injections and echocardiogram measurements. Echocardiogram shows Left Ventricular Ejection Fraction (LVEF) and Right Ventricle (RV) Diameter between different genotypes at baseline (Cre control, n=7; Pkp2-eKO, n=16; Pkp2-cKO, n=14; Pkp2-ceKO, n=9), 21 (Cre control, n=14; Pkp2-eKO, n=19; Pkp2-cKO, n=13; Pkp2-ceKO, n=17) and 42-days dpi (Cre control, n=5; Pkp2-eKO, n=5; Pkp2-cKO, n=6; Pkp2-ceKO, n=5). LVEF and RV Diameter are represented by mean +/- SD. Data analyzed by Two-way ANOVA with Tukey post hoc test. *$P$<0.05 **$P$<0.01, ***$P$<0.001, ****$P$<0.0001. ns, not significant.

**Supplementary Figure 2.** QC plot of scRNA-seq (nCount, nFeature, and percent mitochondrial) between Cre control, Pkp2-cKO, and Pkp2-ceKO samples for scRNA-seq.

**Supplementary Figure 3. A)** Violin plots of vascular endothelial cell markers (*Pecam1, Vwf, Selplg,* and *Icam1*). **B)** Violin plots of lymphatic endothelial cell markers (*Lyve1, Prox1*, and *Flt4*). **C)** Upregulated and Downregulated transcription factor programs in the Pkp2-ceKO mice compared to Pkp2-cKO and Cre control.

**Supplementary Figure 4.** Pseudotime trajectory score and Enrichr analysis of the different fibroblast populations reveals other terminal points in scRNA-seq. Quiescent Fibroblasts (Cluster 0, Lineage 1), Activated Fibroblasts (Cluster 5, Lineage 3), and Pericytes (Cluster 8, Lineage 4).

**Supplementary Figure 5.** UMAP of senescence associated secretory phenotype in fibroblast subsets (*Cxcl14, Cxcl12, Hp, Igfbp4, Cd302, Gas6, Trf, Serping1,* and *Tmem176a*)

**Supplementary Figure 6. A)** UMAP plots of general macrophage/monocyte markers (*Cd86, Mrc1, Itgam, Fcgr1, Timd4, Cd163, Ccl24,* and *Plac8*). **B)** UMAP plots of B cell markers (*Ms4a1, Cr2, Fcer2a, Cd38, Sdc1, Cd27, Fas, Cd69, Cd1d1, Cd27*, and *Cd5*). **C)** UMAP plots of T cell markers (*Cxcr3, Cxcr4, Cxcr5, Il7r, Ifng, Ccr7, Ccr4, Il2ra, Il7r, Foxp3, Stat5a, Sell, Kirb1c, Tnf, Ctla4, Havcr2, Pdcd1,* and *Tnf*).

**Supplementary Figure 7.** CellChat defines intercellular communication **A)** from Pkp2-cKO bulkRNA-seq to pro-inflammatory fibroblasts (top) and **B)** from pro-inflammatory fibroblasts to Pkp2-cKO bulkRNA-seq (bottom).
